# Low-Strength Type I Interferon Signaling Promotes CAR T-Cell Treatment Efficacy

**DOI:** 10.1101/2025.05.13.653878

**Authors:** Erting Tang, Yifei Hu, Guoshuai Cao, Duy-Thuc Nguyen, Nicholas W. Asby, Nada S. Aboelella, Hanna A. Ruiz, Xiaolei Cai, Wenbo Zhang, Yu Zhao, Lishi Xie, Xiufen Chen, Michael R. Bishop, Peter A. Riedell, James L. LaBelle, Justin P. Kline, Jun Huang

## Abstract

CD19-directed chimeric antigen receptor (CAR) T-cell therapy has significantly advanced the treatment landscape for relapsed/refractory diffuse large B-cell lymphoma (r/r DLBCL). However, up to 60% of patients do not achieve a complete response. To uncover determinants of therapeutic efficacy, we analyzed the infusion products of eight r/r DLBCL patients with distinct clinical responses to axicabtagene ciloleucel using single-cell transcriptomics. Compared to patients who exhibited progressive disease, infusion products of complete responders demonstrated enriched signatures of type I interferon (IFN-I) signaling. Based on these findings, we developed a novel strategy to improve CD19-directed CAR T-cell treatment efficacy by incorporating IFN-I as an enhancer during the *ex vivo* manufacturing process, with IFN-I removal before CAR T-cell infusion to avoid *in vivo* toxicities. For both CD28- and 4-1BB-costimulated second-generation CARs, we found that low-strength IFN-I signaling enhanced CAR T-cell cytotoxicity and treatment efficacy against B-cell lymphoma and leukemia. Our low-strength IFN-I-enhanced CAR T-cell *ex vivo* manufacturing approach leverages an existing FDA-approved pharmacologic agent, circumvents *in vivo* interferon-associated toxicities, and remains fully compatible with current CAR constructs and manufacturing workflows. Together, our results establish IFN-I as a potent and costimulation-independent enhancer of CAR T-cell efficacy and provide a translationally feasible approach to enhance CAR T-cell therapies.

## INTRODUCTION

Chimeric antigen receptor (CAR) T-cell therapy has advanced the standard-of-care for hematologic malignancies, achieving complete response rates ranging from 39-71% across relapsed/refractory diffuse large B-cell lymphoma (r/r DLBCL)^1^, acute lymphoblastic leukemia^2^, and multiple myeloma^3,4^. However, the therapy’s effectiveness is constrained by significant rates of treatment failure. In r/r DLBCL, only ∼40% of patients exhibit a complete response, whereas the remaining ∼60% of patients exhibit a partial response, stable disease, or progressive disease, as defined by the International Working Group response criteria.^1^ Patients who cannot clear their lymphoma burden have limited therapeutic options and often succumb to their disease.

Existing studies have correlated treatment failure with diminished CAR T-cell polyfunctionality^5^, increased T-cell exhaustion signatures within CAR T-cell products^6–8^, inhibition by CAR Tregs^9^, as well as failure of CAR T cells to persist and expand *in vivo*^1,10^, among others^11^. Given the importance of CAR T-cell-intrinsic factors in determining treatment outcomes, we investigated the infusion products (IP) of CAR T-cell therapy patients with r/r DLBCL using single-cell transcriptomics. We observed that, compared to patients with progressive disease, the IPs of complete responders demonstrated enriched signatures of type I interferon (IFN-I) signaling. This observation was also validated using an independent, published dataset from Maus et al.^12^

As potent immunomodulators, IFN-I cytokines have long been of interest for their potential to modulate immunotherapy response rates.^13,14^ Upon binding to type I interferon receptors (IFNAR), IFN-I cytokines trigger a phosphorylation cascade that activates transcription factors such as signal transducer and activator of transcription 1 (STAT1), STAT2, and interferon regulatory factor 9 (IRF9).^13^ These transcription factors coordinate transcription of interferon-stimulated genes, such as *IRF7*, *MX1*, *ISG15*, *ISG20*, *IFITM1/2/3*, which mediate antiviral immunity and effector functions.^13,15^ In CD8^+^ T cells, IFN-I signaling enhances cytotoxicity^16^ and induces central memory differentiation^17^, while also controlling T-cell exhaustion^18^, inhibitory receptor expression^19^, and apoptosis^20,21^. Due to their immunomodulatory effects, clinical formulations of IFN-I cytokines (e.g., Avonex, Intron A, Alferon N) have been approved by the FDA for the treatment of malignancies (e.g., hairy cell leukemia, follicular lymphoma)^22^, autoimmune diseases (e.g., multiple sclerosis)^23^, and viral infections (e.g., hepatitis B and C)^24^. Despite being an attractive FDA-approved cytokine for regulating immune functions in cancer, autoimmunity, and infection, IFN-I has not been utilized to modulate CAR T-cell therapy responses.

Given our bioinformatic discoveries and the literature-established roles of T-cell-intrinsic IFN-I signaling in modulating T-cell differentiation and functionality^13^, we hypothesize that IFN-I signaling enhances CAR T-cell efficacy during the *ex vivo* manufacturing process. In this study, we detail our discovery and development of IFN-I cytokines as an enhancer to partially address CAR T-cell therapy failure rates. Through *in vitro* functional assays and *in vivo* lymphoma and leukemia xenograft mouse models, we demonstrate proof-of-principle for a better manufacturing method that leverages low-strength, *ex vivo*, and FDA-approved IFN-I cytokines to improve the *in vivo* treatment efficacy of CAR T-cell therapy.

## MATERIALS AND METHODS

### Collection of patient biospecimens

Deidentified patient biospecimens were gathered in accordance with ethical guidelines under the institutional review board at the University of Chicago Medicine. Informed consent was obtained from all patients or their legal guardians prior to sample collection, in compliance with institutional and federal guidelines. All patients received commercially manufactured axi-cel (Yescarta^®^, Kite Pharma), generated using the same standardized GMP manufacturing protocol at a single external manufacturing facility. Patients were treated at UChicago Medicine between 2019 and 2021 and were selected based on the availability of a clearly defined clinical response, as determined by PET/CT imaging at 1 month post-CAR T-cell infusion, as well as the availability of residual infusion product cells for downstream analyses. Residual infusion product cells were collected from each patient’s infusion bag, stored in CELLBANKER 1 (Amsbio, 11888), and subsequently cryopreserved in liquid-phase nitrogen for subsequent single-cell sequencing.

### Single-cell RNA-seq data generation

Cryopreserved biospecimens were thawed and washed with cold FACS buffer (PBS, 2% BSA, 0.05% sodium azide). Cells were conjugated with LIVE/DEAD Fixable Near-IR viability dye (Invitrogen, L34975) at 1:1000 dilution in PBS for 5 minutes at room temperature. Finally, cells were washed 3x in cold cell media (RPMI, 10% FBS) before sorting (BD Biosciences, FACSAria Fusion) for live cells. Sorted cells were partitioned into droplets for single-cell RNA-seq via Chromium Next GEM Single-Cell 5’ Kit v2 (10x Genomics, 1000263). RNA-seq libraries were prepared according to manufacturer protocols. Libraries were quantified via the Qubit dsDNA HS Assay Kit (Invitrogen, Q32851), quality-checked for fragment sizes via high-sensitivity D5000 screentapes (Agilent, 5067-5592), pooled, and sequenced (Illumina, Novaseq-6000, RRID:SCR_016387 and Nextseq-550, RRID:SCR_016381).

### Single-cell RNA-seq data analysis

Alignment, filtering, counting, and demultiplexing of RNA-seq reads were performed using the Cell Ranger *count* (10x Genomics, version 7.1.0, RRID:SCR_017344). For alignment, the GRCh38 reference genome was appended with the CAR transgene sequence of axicabtagene ciloleucel.

Downstream analysis was performed using Seurat^25^ (version 4.3.0, RRID:SCR_016341). After converting count matrices and merging all Seurat objects, we filtered cells with >15% mitochondrial reads and 40,000 UMIs to remove debris, doublets, and dead cells. The matrix was normalized using *NormalizeData* with default options. The top 5,000 variable genes found by *FindVariableFeatures* with the default “vst” method were taken as variable genes. Data were centered and scaled using *ScaleData*, with additional regression against (1) the percent of mitochondrial RNA content and (2) difference in cell cycle S-phase and G2M-phase scores. Scaled data were then used for principal component analysis (PCA) using *RunPCA*. Data harmonization to remove patient-specific effects was performed on the principal components using Harmony^26^ (version 0.1.1, RRID:SCR_018809) through *RunHarmony*. Top 50 Harmony components were used for Uniform Manifold Approximation and Projection (UMAP) dimensionality reduction via *RunUMAP*. The same harmony components were used to construct the shared nearest neighbor (SNN) graph using *FindNeighbors*, which was then partitioned to identify clusters using *FindClusters* with the “original Louvain” algorithm.

Differentially expressed genes were determined using *FindMarkers* in the Seurat package with default parameters. Gene set enrichment analysis and pathway enrichment analysis were carried out using clusterProfiler^27^ (version 4.6.2, RRID:SCR_016884) based on pathways from the msigdbr database’s built-in msigdbr^28^ package (version 7.5.1, RRID:SCR_022870). Gene module scores were calculated for each cell using *AddModuleScore* in Seurat for msigdbr or customized gene sets.

### Cell lines

The OCI-Ly8 (RRID:CVCL_8803) cell line was obtained from Dr. James L. LaBelle’s lab (University of Chicago), the VAL cell line (RRID:CVCL_1819) was obtained from Dr. Justin P. Kline’s lab (University of Chicago), the NALM6 cell line was obtained from ATCC (CRL-3273™, RRID:CVCL_0092), and the Lenti-X 293T packaging cell line was obtained from Takara (632180, RRID:CVCL_4401). All cell lines were confirmed to be mycoplasma-negative using the Universal Mycoplasma Detection Kit (ATCC, 30-1012k), with the most recent testing performed in November 2025. All cell lines were used for experiments within three passages after thawing.

### Lentiviral production and transduction

CD28- or 4-1BB-costimulated anti-CD19 human CARs were designed based on the sequences of axi-cel or liso-cel. The axi-cel sequence was obtained from a previously published sequencing study,^29^ while the liso-cel sequence was retrieved from the Global Substance Registration System database (UNII: 7K2YOJ14X0). A GFP-tagged, CD28-costimulated or 4-1BB-costimulated, anti-CD19 human CAR was cloned onto the pHR vector backbone to constitute the transfer plasmid. Plasmids were transfected into the Lenti-X 293T packaging cell line (Takara, 632180, RRID:CVCL_4401) for lentiviral production. In brief, Lenti-X 293T cells were seeded and grown overnight to ∼90% confluence. Subsequently, transfer, packaging (psPAX2, Addgene, #12260, RRID:Addgene_12260), and envelope (pMD2.G, Addgene, #12259, RRID:Addgene_12259) plasmids were co-transfected into Lenti-X 293T cells via Lipofectamine 3000 (Invitrogen, L3000001). After 72 hours, the viral supernatant was collected and stored at -80°C until transduction. During transduction, lentiviruses were added to human primary T cells along with protamine sulfate (Millipore Sigma, P3369-10G) to a final concentration of 10 µg/mL. Cells were spinoculated at 800 × *g* for 60 minutes at room temperature. All cell lines were routinely tested for Mycoplasma contamination every 4-6 weeks using a PCR-based assay.

### CAR T-cell production and culturing

PBMC samples were collected from healthy donors under an Institutional Review Board approved protocol. Cryopreserved human PBMCs were thawed in warm T-cell media (RPMI supplemented with 10% FBS, 1% penicillin/streptomycin, 2 mM L-glutamine, and 50 μM 2-mercaptoethanol). Subsequently, PBMCs were cultured for 3 hours in complete T-cell media (T-cell media supplemented with 50 units/mL of IL-2) in order to allow monocytes to adhere to the culture flask. The non-adhered cells were transferred to a new culture flask. Cells were counted via the LUNA-FL cell counter. T-cell percentage (% CD3^+^) was calculated by flow cytometry.

T cells were seeded in complete T-cell media at 0.75 × 10^6^/mL. CD3/CD28 Dynabeads (Gibco, 11161D) were washed and added at 1:1 ratio. After overnight activation, T cells were transduced with CAR lentiviruses (MOI of 5). The media was refreshed after 24 hours with complete T-cell media dosed with recombinant human IFN-α2 (BioLegend, 592702). T cells were subsequently expanded, and media was refreshed every 2 days.

### Quantitative reverse transcription PCR

CAR T cells were treated under IFN-I culturing conditions for 24 hours. Then, cells were collected, and stained for viability (Invitrogen, L34975). Live CAR T cells were sorted (BD Biosciences, FACSAria Fusion). Total RNA was extracted using the Monarch Total RNA Miniprep Kit (NEB, T2010S), according to manufacturer protocols. RNA was quantified via the Qubit RNA HS Assay Kit (Invitrogen, Q32852). cDNA was synthesized using the GoScript Reverse Transcription System (Promega, A5000).

Quantitative PCR was performed using Taqman Gene Expression Assay (ThermoFisher Scientific, 4331182) and Applied Biosystems TaqMan Universal PCR Master Mix (ThermoFisher Scientific, 4304437), on LightCycler® 96 Instrument (Roche). The cycling conditions were as follows: UNG incubation at 50°C for 2 minutes, initial denaturation at 95°C for 10 minutes, followed by 40 cycles of denaturation at 95°C for 15 seconds and annealing/extension at 60°C for 60 seconds. Each sample was run in triplicate.

The specific Taqman Gene Expression Assay used were as follows: *ACTB* (Hs01060665_g1), *IRF7* (Hs01014809_g1), *MX1* (Hs00895608_m1), and *ISG15* (Hs01921425_s1). Relative gene expression levels were calculated using the 2^-ΔΔCt method, with *ACTB* serving as the internal control. Statistical analysis was performed using one-way ANOVA followed by a Tukey’s post hoc test for significance.

### In vitro co-culture immunoassays

For all co-culture immunoassays, CAR T cells (E) and luciferized OCI-Ly8 cells (T, RRID:CVCL_8803) were seeded under a fixed E/T ratio. To measure *in vitro* cytotoxicity, co-cultures were established with E/T ratio of 1:1. After 6 hours, One-Glo Luciferase substrate (Promega, E6110) was added to each well. Residual tumor cells were quantified via luciferase activity. As a positive control, target cells were directly lysed with 1% NP-40 (Abcam, ab142227). To measure cytotoxic molecule expression, co-cultures were established with E/T ratio of 1:1 with 1× brefeldin A (BioLegend, 420601). After 4 hours, cells were collected for intracellular staining and flow cytometry. To measure cytokine secretion, co-cultures were established with E/T ratio of 1:1. After 24 hours, the culture supernatant was collected and stored. Secreted cytokines were quantified using LEGENDplex immunoassays per manufacturer’s protocol (BioLegend). To measure *in vitro* persistence, co-cultures were established with E/T ratio of 1:5. Every three days, cells were counted using Precision Count Beads (BioLegend, 424902) and measured for apoptosis on flow cytometry. Cell media was then refreshed.

### Flow cytometry

Cells were washed with cold FACS buffer (PBS, 2% BSA, 0.05% sodium azide). Fc receptors were blocked by incubation with Human TruStain FcX (BioLegend, 422301) at 1:50 dilution for 5 minutes at 4°C. Then, cells were incubated for 30 minutes at 4°C in the dark with an antibody staining solution for surface antigen staining. Antibodies were used according to manufacturer recommendations. After staining, cells were incubated briefly either with live/dead staining solution or apoptosis staining solution for 10 minutes at room temperature. Then, cells were either washed and resuspended in FACS buffer or resuspended in Annexin V Binding Buffer (BioLegend, 422201). Cells were ready for analysis by flow cytometry or intracellular staining.

For intracellular staining, cells stained for surface antigens were subject to the BD Fixation/Permeabilization Kit (BD Biosciences, 554714) for cytotoxic molecule staining or True-Nuclear™ Transcription Factor Buffer Set (BioLegend, 424401) for IRF7 staining, following manufacturer recommendations. In brief, cells were fixed in fixation buffer, washed 3× in permeabilization buffer, and incubated for 60 minutes at 4°C in the dark with a staining solution containing antibodies and permeabilization buffer. After staining, cells were washed 2× in permeabilization buffer, 1x in FACS buffer, and resuspended in FACS buffer for analysis by flow cytometry.

For Ki-67 staining, cells stained for surface antigens were washed 2× in PBS, fixed in prechilled 70% ethanol overnight, washed 2× in FACS buffer, and incubated for 30 minutes at 4°C in the dark with Ki-67 antibody staining solution. After staining, cells were washed 2× in FACS buffer, and resuspended in FACS buffer for analysis by flow cytometry.

Flow cytometry was performed on LSRFortessa (BD Biosciences, RRID:SCR_018655) or Aurora (Cytek Biosciences, RRID:SCR_019826). Data were analyzed using FlowJo software (version 10.10.0, RRID:SCR_008520).

### Antibodies

All antibodies were used according to manufacturer recommendations. APC-anti-IFNAR2 (10359-MM07T-A) was from Sino Biological. All other antibodies originate from BioLegend. For surface staining, AF647-anti-CD3 (clone: UCHT1, 300416, RRID:AB_389332), BV421-anti-CD3 (clone: UCHT1, 300434, RRID:AB_10962690), BV510-anti-CD4 (clone SK3, 344633, RRID:AB_2566016), PerCP/Cy5.5-anti-CD8α (clone SK1, 344709, RRID:AB_2044009), PE-anti-CD45RA (clone HI100, 304107, RRID:AB_314411), BV421-anti-CD45RO (clone UCHL1, 304223, RRID:AB_10898323), AF647-anti-CCR7 (clone G043H7, 353217, RRID:AB_10913812), BV421-anti-PD-1 (clone EH12.2H7, 329919, RRID:AB_10900818), PE/Cy7-anti-LAG-3 (clone 11C3C65, 369309, RRID:AB_2629752), PE/Dazzle™ 594-anti-TIGIT (clone A15153G, 372715, RRID:AB_2632930), PE/Cy7-anti-CCR7 (clone G043H7, 353225, RRID:AB_11125576), and BV785-anti-CD62L (clone DREG-56, 304829, RRID:AB_2629516) were used. For intracellular staining, PE/Cy7-anti-granzyme B (clone QA16A02, 372213, RRID:AB_2728380), PE-anti-perforin (clone B-D48, 353303, RRID:AB_10915476), and PE-anti-IRF7 (clone 12G9A36, 656003, RRID:AB_2563006) were used. For live/dead staining, Zombie NIR Fixable Viability Kit (423106) was used in PBS. For apoptosis staining, BV421-Annexin V (640923, RRID:AB_2893503) or PE/Cy7-Annexin V (640950) was diluted in Annexin V Binding Buffer (422201). For Ki-67 staining, PE-anti-Ki-67 (clone Ki-67, 350503, RRID:AB_10660818) was used.

### Xenograft mouse models

Immunocompromised NOD-*scid* IL2Rgamma^null^ (NSG, Strain #005557) mice were purchased from JAX and maintained at the animal resource center (ARC) of the University of Chicago in accordance with institutional guidelines. Mice were monitored daily. Care and experimental procedures were carried out in line with the Institutional Animal Care and Use Committee (IACUC) approved protocols of the Animal Care Center at the University of Chicago.

For the standard xenograft models, 5×10^6^ OCI-Ly8-Chili-luc or 5×10^5^ NALM6-Chili-luc cells in 100 µL PBS were intravenously injected into 8-12-week-old NSG mice. Mice were randomized prior to CAR T-cell injection. 5×10^5^ or 1×10^6^ CAR T cells were intravenously injected 1 week later for the OCI-Ly8 model or 4 days later for the NALM6 model. For the OCI-Ly8 model to evaluate durable efficacy of CAR T cells, 3×10^6^ OCI-Ly8-Chili-luc cells were intravenously injected on day 0, followed by randomization and intravenous injection of 3×10^5^ CAR T cells 1 week later. Mice were then injected with an additional 3×10^6^ OCI-Ly8-Chili-luc cells intravenously 10 days after CAR T-cell infusion. Tumor progression was monitored using either IVIS Spectrum (PerkinElmer) or Lago X (Spectral Instruments Imaging) following the intraperitoneal injection of 1 mg D-luciferin potassium (PerkinElmer, 122799) in 200 µL PBS. Animals were anesthetized at 1.5% isoflurane throughout acquisition. Bioluminescence was quantified with Aura software (Spectral Instruments Imaging). Euthanasia was conducted via CO_2_ overdose followed by cervical dislocation upon manifestation of paralysis, impaired mobility, or poor body condition (BCS<2). Group sizes were determined based on prior experience with the xenograft model and expected treatment effect sizes.

Blood was collected by cheek bleeding using a 4-mm lancet (Fisher Scientific, NC9922361), and 50 µL was subjected to red blood cell (RBC) lysis using RBC lysis buffer (eBioscience, 00-4300-54). Lysis was quenched with FACS buffer, and cells were washed once with FACS buffer before proceeding to flow staining. Spleens were collected by surgery and gently ground through a 40-µm cell strainer, followed by RBC lysis using RBC lysis buffer (eBioscience, 00-4300-54). After stopping the lysis with RPMI supplemented with 10% FBS, cells were washed once more with RPMI + 10% FBS before being subjected to flow staining.

### Statistical analysis

Statistical tests were performed on R (version 4.2.1, RRID:SCR_001905) or GraphPad Prism 10 (version 10.1.1, RRID:SCR_002798). Comparisons between two groups were performed using Wilcoxon rank-sum test or unpaired t-test. Comparisons between more than two groups were performed by either one-way or two-way ANOVA followed by Tukey’s correction for multiple comparisons. Survival data were analyzed using the Log-Rank (Mantel-Cox) test.

### Data availability

All data are available from the authors upon reasonable request.

### Code availability

Codes supporting the analyses performed in this study are available at https://github.com/ertingtang/IFN-I-CART.

## RESULTS

### Enrichment of IFN-I signaling in infusion products of complete responders

To uncover determinants of therapeutic efficacy, infusion products (IPs) from 8 r/r DLBCL patients treated with CD28-costimulated CD19-directed CAR T-cell therapy (axicabtagene ciloleucel, “axi-cel”) at the University of Chicago Medicine were profiled using single-cell sequencing. The cohort included 5 patients who achieved a complete response (CR) at one month post-infusion and another 3 patients who exhibited progressive disease (N-CR) (**Fig. 1A**, Supplementary Table 1). After removing low-quality cells, we obtained 27,879 cells distributed among 7 T-cell clusters (**Fig. 1B**) based on gene expression (**Fig. 1C**, Supplementary Fig. S1A). No cluster was patient specific (**Fig. 1D**, Supplementary Fig. S1B). The two CD8^+^ clusters were annotated as central memory (CM, *SELL*^+^*IL7R*^+^*TBX21^−^KLRG1^−^*) or effector memory (EM, *SELL^−^IL7R*^+^*TBX21*^+^*KLRG1*^+^) T cells. The two CD4^+^ clusters were annotated as memory (Mem, *FOXP3*^lo^*IL7R*^+^*CTLA4*^lo^) or regulatory (Treg, *FOXP3*^hi^*IL7R^−^CTLA4*^hi^) T cells. The remaining three clusters consisted of naïve-activated (*SELL*^+^*IL7R*^+^*TCF7*^+^*CD38*^+^), unconventional (*TRDC*^+^*NCAM1*^+^) as a mixture of γδ and natural killer phenotypes, and proliferating (*MKI67*^+^*TOP2A*^+^) T cells. No discernable composition differences were found between CR and N-CR patients (**Fig. 1B** and **D**, Supplementary Fig. S1C).

**Figure 1.**
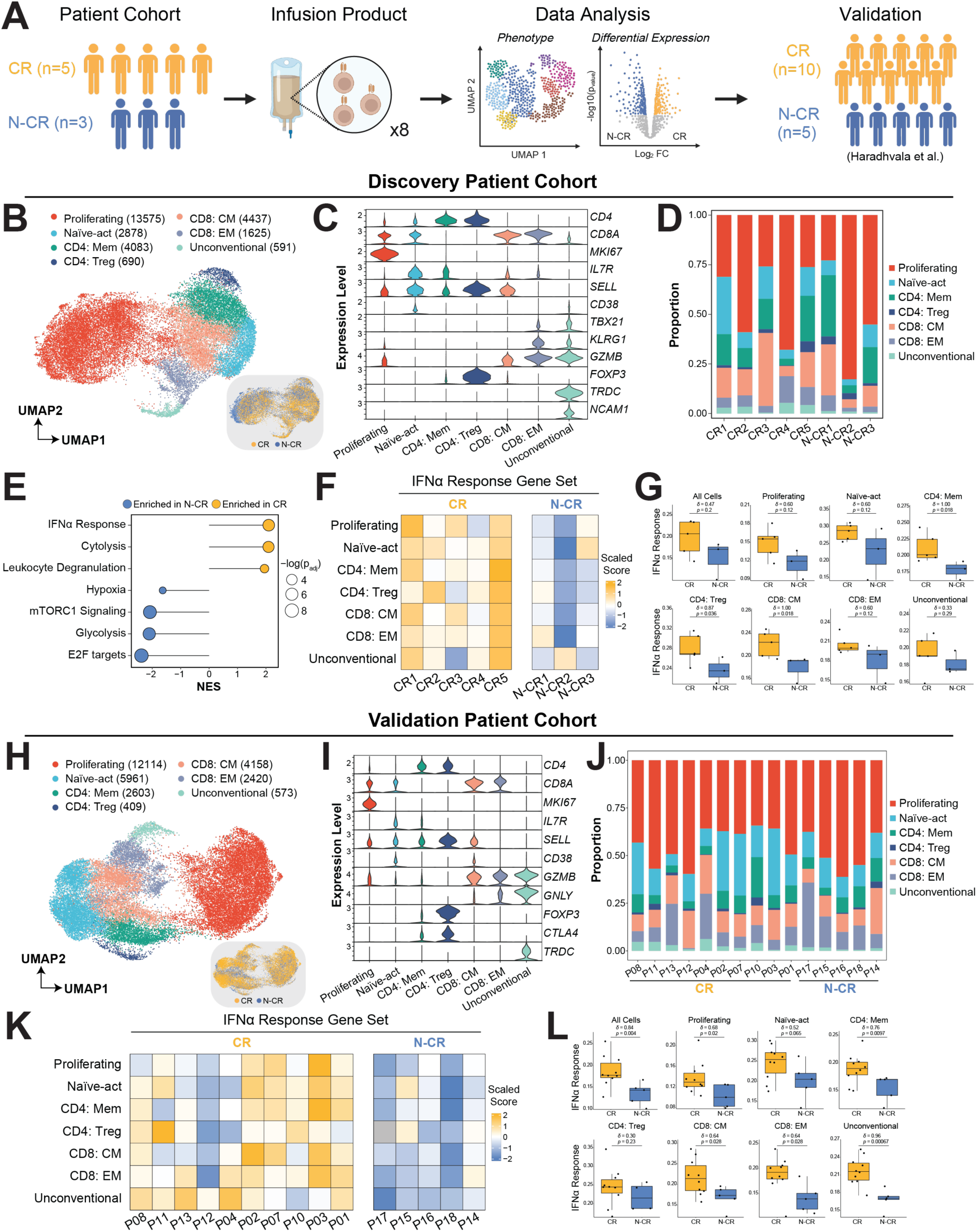
Enrichment of IFN-I signaling in infusion products of complete responders. **A,** Cartoon depicting study design of single-cell analysis on infusion products of patients who received axi-cel from discovery and validation datasets. CR, complete responder, N-CR, non-complete responder. **B,** UMAP demonstrating 7 T-cell clusters (proliferating, Naïve-activated, CD4^+^ Memory, Treg, CD8^+^ central memory, CD8^+^ effector memory, unconventional) in infusion products from discovery dataset. No patient-specific or response-specific clusters were found. **C,** Violin plots depicting normalized expression levels of representative markers for each cluster. **D,** Stacked bar graphs depicting T-cell cluster proportions for each patient in discovery dataset. **E,** Gene set enrichment analysis comparing CR and N-CR T cells. For each gene set, direction and statistical significance of enrichment are indicated by the circle’s color and size, respectively. **F,** Tilemap depicting scaled IFNα response signature scores in each T-cell cluster across patients. **G,** Box plots depicting IFNα-response signature scores for CR and N-CR T cells within each T-cell cluster. Each dot represents one patient. Effect sizes were calculated using Cliff’s δ, and statistical significance was assessed using the Wilcoxon rank-sum test. **H,** UMAP demonstrating 7 T-cell clusters in infusion products from validation dataset from Maus et al. **I,** Violin plots depicting normalized expression levels of representative markers for each cluster. **J,** Stacked bar graphs depicting T-cell cluster proportions for each patient in validation dataset. **K,** Tilemap depicting scaled signature scores of IFNα response gene sets in each T-cell cluster across patients. No Treg was found in P17 infusion products, colored in gray. **L,** Box plots depicting IFNα-response signature scores for CR and N-CR T cells within each T-cell cluster. Each dot represents one patient. Effect sizes were calculated using Cliff’s δ, and statistical significance was assessed using the Wilcoxon rank-sum test.

Gene set enrichment analysis revealed distinct differences between CR and N-CR T cells. CR T cells were enriched for signatures of type I interferon signaling (“IFNα response”), cytolysis, and leukocyte degranulation, whereas N-CR T cells were enriched for signatures of cell growth, hypoxia, and glycolysis (**Fig. 1E**). Notably, the CR-specific IFNα response signature was enriched in CR T cells across T-cell clusters and individual patients, with the effect-size analysis (Cliff’s δ) indicating moderate to strong differences between CR and N-CR groups (**Fig. 1F** and **G**).

To validate our discovery of CR-specific IFN-I signaling signature, we tested our findings using an independent and published single-cell transcriptomic dataset from Maus et al. (*n* = 15 patients with lymphoma who received axi-cel, **Fig. 1A**).^12^ This cohort included 10 patients who achieved a complete response (CR) and 5 who did not (N-CR), as assessed one-month post-infusion. After filtering out low-quality cells, performing dimensionality reduction, and clustering, we uncovered the same 7 T-cell clusters identified in our own dataset (**Fig. 1H**) based on gene expression (**Fig. 1I**, Supplementary Fig. S2A). No cluster was patient specific (**Fig. 1J**). Notably, expression of ISGs (including *ISG20*, *IFITM1/3*) was upregulated in IPs of CR (Supplementary Fig. S2B). Most importantly, IFNα response signature was enriched in IPs of CR (Supplementary Fig. S2C). The CR-specific enrichment of IFNα response signature was consistently observed across all clusters (except Tregs) and across individual patients, again with effect-size analysis (Cliff’s δ) indicating moderate to strong differences between CR and N-CR groups (**Fig. 1K** and **L**).

Taken together, findings from the discovery and validation datasets strongly support an association between IFN-I signaling and improved CAR T-cell clinical efficacy.

### IFN-I signaling increases CAR T-cell cytotoxicity

Previous studies have demonstrated the immunomodulatory role of T cell-intrinsic IFN-I signaling.^13^ For instance, IFN-I signaling promotes cytotoxicity^16^, T_h_1 polarization^30–32^, and central memory phenotype differentiation^17^, all of which may be desired CAR T-cell characteristics and potentially explain the differential clinical responses observed in the two patient cohorts. Based on these findings and our bioinformatics analyses, we hypothesized that IFN-I signaling can enhance CAR T-cell efficacy. To test this hypothesis, we generated both CD28- and 4-1BB-costimulated CD19-directed CAR T cells using primary T cells from healthy human donors, based on axicabtagene ciloleucel (axi-cel, **Fig. 2B**) and lisocabtagene maraleucel (liso-cel, **Fig. 2C**) respectively. To induce IFN-I signaling, we treated CAR T cells with exogenous IFN-α2, a highly potent, FDA-approved, and clinically compatible IFN-I subtype, for culturing CAR T cells (**Fig. 2A**). Prior to addition of IFN-α2, we confirmed that CAR T cells globally expressed the type I interferon receptor in a transduction-independent and costimulation-independent manner (Supplementary Fig. S3A and C).

**Figure 2.**
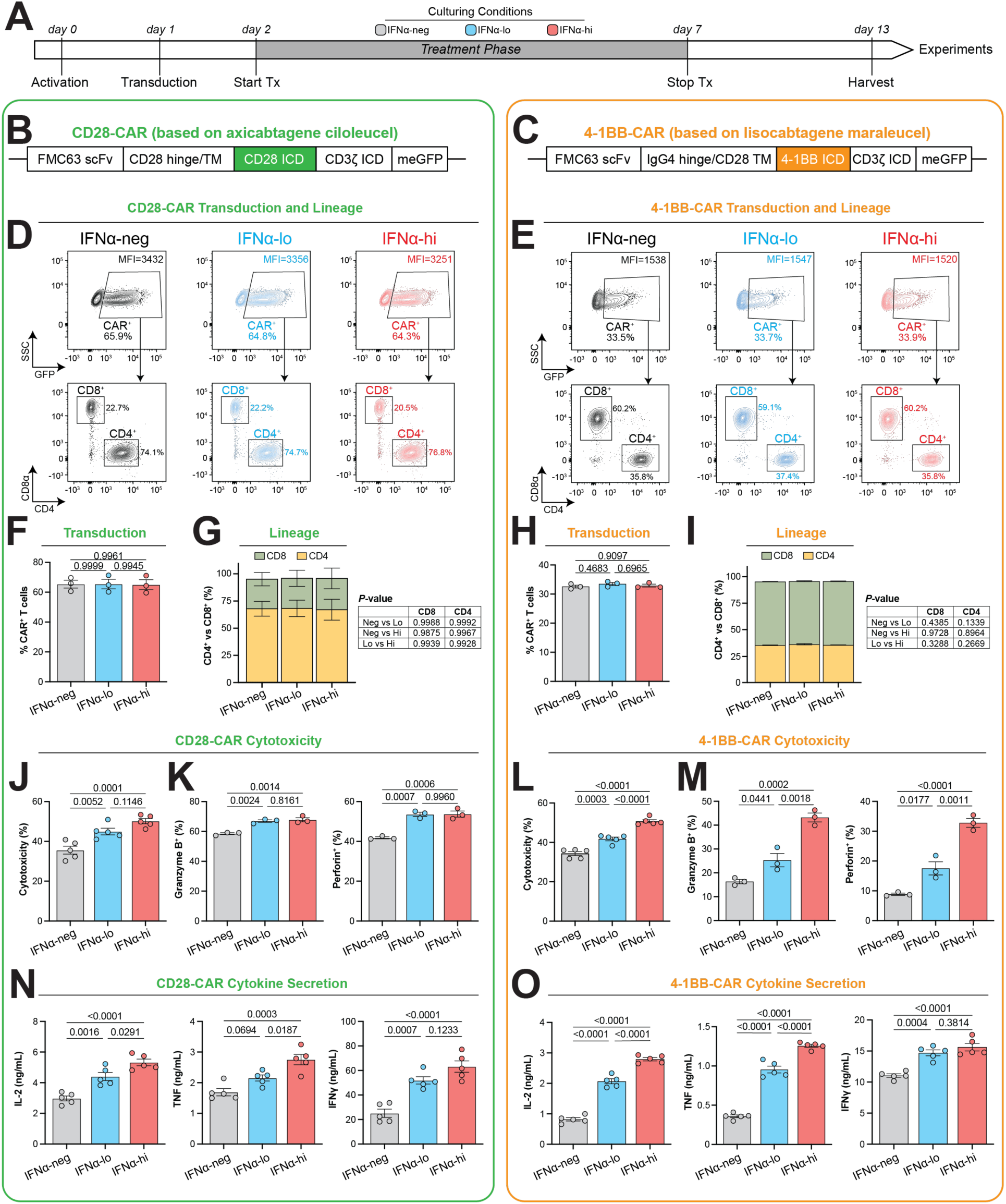
IFN-I signaling increases CAR T-cell cytotoxicity. **A,** Workflow of CAR T-cell *ex vivo* generation, treatment, and culturing. Both CD28- (axi-cel-based) and 4-1BB-costimulated (liso-cel-based) CAR T cells were generated for analysis. **B** and **C,** CD28- or 4-1BB-costimulated CAR designs based on the construct used in axicabtagene ciloleucel (**B**) or lisocabtagene maraleucel, respectively (**C**). **D** and **E,** Representative flow plots depicting gating for CAR expression (top) and T-cell lineages (bottom) in CD28-costimulated (**D**) and 4-1BB-costimulated (**E**) CAR T cells. CAR T cells were gated as live CD3^+^meGFP^+^ cells. **F** and **H,** Transduction percentage as measured by meGFP fluorescence among T cells from each culturing condition in CD28-costimulated (**F**, n=3) and 4-1BB-costimulated (**H**, n=3) CAR T cells. **G** and **I,** Stacked bar plots comparing CD8 vs CD4 proportions in CD28-costimulated (**G**, n=3) and 4-1BB-costimulated (**I**, n=3) CAR T cells. **J**-**M,** Cytotoxicity of CD28-costimulated (**J**-**K**) and 4-1BB-costimulated (**L**-**M**) CAR T cells. **J** and **L,** Direct cytotoxicity against firefly luciferase-expressing OCI-Ly8 target cells at E/T ratio of 1:1 via 6-hr bioluminescence assay (n=5). **K** and **M,** Intracellular staining for granzyme B (right) and perforin (left) after 4-hr of coculture with OCI-Ly8 at E/T ratio of 1:1 as measured by flow cytometry (n=3). **N** and **O,** Concentration of cytokines secreted by CD28-costimulated (**N**) and 4-1BB-costimulated (**O**) CAR T cells into the supernatant after 24-hr of coculture with OCI-Ly8 at E/T ratio of 1:1 as measured by ELISA (n=5). From left to right, IL-2, TNF, and IFNγ. All data are shown as mean ± SEM of n experimental replicates from a representative donor. Results are representative of independent experiments using CAR T cells from 3 different donors. Statistical analysis were performed via one-way (**F**, **H**, **J**-**O**) or two-way (**G** and **I**) ANOVA with Tukey’s correction for multiple comparisons.

Since each downstream target of IFN-I signaling exhibits varying sensitivity to IFN-I concentrations^33^, we hypothesized that the IFN-I culturing concentration may affect CAR T-cell functions. To determine the IFN-I concentrations for our study, we initially titrated IFN-α2 on both CD28- and 4-1BB-costimulated CAR T cells and measured intracellular interferon regulatory factor 7 (IRF7), a key ISG^34^ (Supplementary Fig. S3B and D). IFN-I signaling reached saturation at concentrations above 3000 IU/mL of IFN-α2, consistent with previous studies.^16,32^ We also observed half-maximal IRF7 upregulation at approximately 100-300 IU/mL of IFN-α2. Consequently, we established three culturing conditions: IFN-α2 at saturating concentration (“IFNα-hi”, 3000 IU/mL), IFN-α2 near half-maximal effective concentration (“IFNα-lo”, 300 IU/mL), and no IFN-α2 (“IFNα-neg”, 0 IU/mL). We confirmed that these conditions resulted in concentration-dependent upregulation of ISGs, including *MX1*, *IRF7*, and *ISG15* (Supplementary Fig. S4).

CAR T cells were treated based on the three culturing conditions for downstream experimentation (**Fig. 2A**). Briefly, T cells were transduced with the CAR construct one day after activation (day 1). IFN-I treatment was administered from day 2 to day 7, after which IFN-I was completely removed by washing and refreshing the culture medium. Cells were subsequently expanded until day 13 for experiments. Since IFN-I signaling can inhibit viral transduction^35^, we chose to add IFN-α2 after CAR transduction, which enabled us to achieve consistent transduction efficiency and expression of both CD28-costimulated (**Fig. 2D** and **F**) and 4-1BB-costimulated CARs (**Fig. 2E** and **H**) across the three culturing conditions. IFN-α2 was also removed on day 7 of *ex vivo* expansion to prevent residual cytokine from inducing potential *in vivo* toxicity.^36,37^ Between culturing conditions, IFN-I signaling did not affect CD4/CD8 ratios (**Fig. 2G** and **I**), memory phenotypes (Supplementary Fig. S5A-D), or IFN-I receptor expression (Supplementary Fig. S5E-H). Although long-term sustained IFN-I signaling drives T cell exhaustion in tumor microenvironment^18^, our short-term IFN-I exposure in *ex vivo* culture does not induce exhaustion (Supplementary Fig. S5I and J). To test cytotoxicity, CAR T cells were co-cultured with OCI-Ly8 target cells, a CD19^+^ human DLBCL cell line. In both CD28- and 4-1BB-costimulated CAR T cells, we observed IFN-I signaling significantly increased CAR T-cell cytotoxicity (**Fig. 2J** and **L**), expression of cytolytic molecules (granzyme B, perforin, **Fig. 2K** and **M**), and secretion of cytokines (IL-2, TNF, IFNγ, **Fig. 2N** and **O**), in a concentration-dependent manner. These findings suggest that IFN-I signaling increases CAR T-cell cytotoxicity.

### High-strength IFN-I signaling increases apoptosis, whereas low-strength IFN-I signaling does not

The IFNα-lo condition consistently yielded comparable levels of viable CAR T cells to IFNα-neg controls following manufacturing, whereas the IFNα-hi condition reduced CAR T-cell yield (**Fig. 3A** and **C**). Given previous reports that IFN-I signaling can inhibit cell proliferation^38^, we evaluated whether CAR T-cell proliferation was affected by IFN-I signaling through assessing the proliferation marker, Ki-67. No significant differences in Ki-67 expression were observed across the three culturing conditions, indicating that IFN-I signaling does not impair CAR T-cell proliferation during *ex vivo* expansion (**Fig. 3B** and **D**; Supplementary Fig. S6). Since IFN-I signaling also reportedly promotes T-cell apoptosis^20,21^, we next quantified CAR T-cell apoptosis via Annexin V staining. Flow phenotyping revealed that CAR T cells in the IFNα-hi condition trended towards increased apoptosis, whereas those in the IFNα-lo condition did not (**Fig. 3E** and **F**).

**Figure 3.**
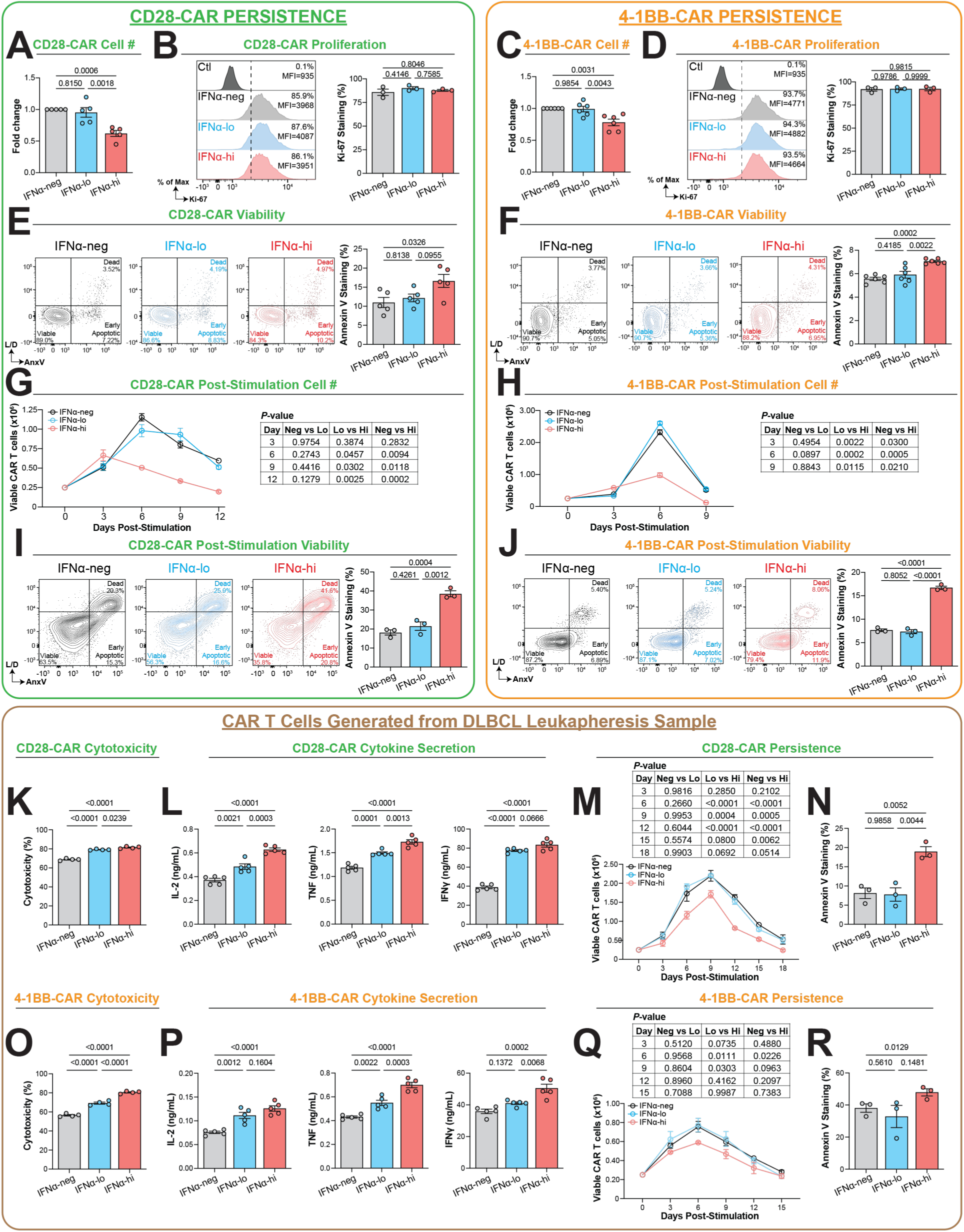
High-strength IFN-I signaling increases apoptosis, whereas low-strength IFN-I signaling does not. **A** and **C,** Relative counts of live CAR T cells from the three culturing conditions after CAR T-cell manufacturing in CD28-costimulated (**A**, n=5) and 4-1BB-costimulated (**C**, n=6) CAR T cells. Counts were normalized to the IFNα-neg condition. **B** and **D,** Proliferation of CAR T cells from the three culturing conditions in CD28-costimulated (**B**, n=3) and 4-1BB-costimulated (**D**, n=3) CAR T cells. Left, representative flow plots depicting gating for Ki-67^+^ proliferating CAR T cells at the day when IFNα was removed. Right, Ki-67^+^ proliferating cell proportions among CAR T cells. Non-activated T cells from human PBMCs were used as a biological control for gating. **E** and **F,** Apoptosis of CAR T cells from the three culturing conditions after CAR T-cell manufacturing in CD28-costimulated (**E**, n=5) and 4-1BB-costimulated (**F**, n=6) CAR T cells. Left, representative flow plots depicting gating for viable (L/D^−^Annexin V^−^), early apoptotic (L/D^−^Annexin V^+^), and dead (L/D^+^Annexin V^+^) cells among CAR T cells. Right, staining for Annexin V positivity among live CAR T cells. **G** and **H,** Line graphs depicting viable CAR T-cell cellularity of CD28-costimulated (**G**, n=3) and 4-1BB-costimulated (**H**, n=3) CAR T cells *in vitro* following coculture with OCI-Ly8 at E/T ratio of 1:5. **I** and **J,** Apoptosis of CD28-costimulated CAR T cells (**I**, n=3) and 4-1BB-costimulated CAR T cells (**J**, n=3) from the three culturing conditions at day 6 after coculture with OCI-Ly8 cells at E/T ratio of 1:5. Left, representative flow plots depicting gating for viable (L/D^−^Annexin V^−^), early apoptotic (L/D^−^Annexin V^+^), and dead (L/D^+^Annexin V^+^) cells among the three culturing conditions. Right, staining for Annexin V positivity among live CAR T cells. **K**-**R,** *In vitro* data of CAR T cells derived from DLBCL patient leukapheresis products. **K** and **O,** Cytotoxicity of CD28-costimulated (**K**, n=4) and 4-1BB-costimulated (**O**, n=4) CAR T cells. **L** and **P,** Concentration of cytokines secreted by CD28-costimulated (**L**, n=5) and 4-1BB-costimulated (**P**, n=5) CAR T cells into the supernatant after 24-hr of coculture with OCI-Ly8 at E/T ratio of 1:1 as measured by ELISA. From left to right, IL-2, TNF, and IFNγ. **M** and **Q,** Line graphs depicting viable CAR T-cell cellularity of CD28-costimulated (**M**, n=3) and 4-1BB-costimulated (**Q**, n=3) CAR T cells *in vitro* following coculture with OCI-Ly8 at E/T ratio of 1:5. **N** and **R,** Apoptosis of CD28-costimulated CAR T cells (**N**, n=3) and 4-1BB-costimulated CAR T cells (**R**, n=3) from the three culturing conditions at the time points when cellularity declined in (**M**) and (**Q**): day 9 for (**N**) and day 6 for (**R**) at an E:T ratio of 1:5. All data are shown as mean ± SEM of n experimental replicates from a representative donor. **A-J** show results representative of independent experiments using CAR T cells from 3 donors. Statistical analysis were performed via one-way (**A**-**F**, **I**-**J**, **K**-**L**, **N-P**, and **R**) or two-way (**G**, **H**, **M**, **Q**) ANOVA with Tukey’s correction for multiple comparisons.

To further investigate the cellularity and apoptosis of CAR T cells from the three culturing conditions following antigen-specific stimulation, we subsequently quantified cellularity and apoptosis of CAR T cells after stimulation with OCI-Ly8 target cells. Following stimulation, CAR T-cell cellularity from the IFNα-lo and IFNα-neg conditions was statistically indistinguishable (**Fig. 3G** and **H**). On the other hand, CAR T-cell cellularity from the IFNα-hi condition was significantly depleted compared to the other two conditions. At the timepoint when cellularity declines became apparent (day 6 post-stimulation in the representative data), we observed a corresponding enrichment in apoptosis in CAR T cells from the IFNα-hi condition, but not in those from the IFNα-lo condition (**Fig. 3I** and **J**).

Taken together, these findings suggest that high-strength, but not low-strength IFN-I signaling increases CAR T-cell apoptosis, thereby negatively affecting persistence during both *ex vivo* expansion and post antigen-specific activation.

### Low-strength IFN-I signaling increases the efficacy of DLBCL patient-derived CAR T cells

To evaluate the clinical translation potential of our IFN-I enhancement process, we tested IFN-α2 with CAR T cells generated from the leukapheresis product of a patient with DLBCL. IFN-I signaling increased CAR T-cell cytotoxicity (**Fig. 3K** and **O**) and cytokine secretion (IL-2, TNF, IFNγ, **Fig. 3L** and **P**) in a dose-dependent manner. Upon stimulation with OCI-Ly8 target cells, CAR T cells from the IFNα-hi condition demonstrated decreased persistence and increased apoptosis, whereas those from the IFNα-lo condition maintained similar persistence compared to IFNα-neg controls without increased apoptosis (**Fig. 3M, N, Q**, and **R**). Findings were consistent across both CD28- and 4-1BB-costimulated CAR constructs (**Fig. 3K-N** compared to **Fig. 3O-R**). These data demonstrate that low-strength IFN-I signaling enhances effector function without inducing apoptosis or compromising persistence of patient-derived CAR T cells. The results also pave the way for future clinical translation of our findings.

### Low-strength IFN-I signaling promotes CAR T-cell treatment efficacy in vivo

Given that the IFNα-lo condition uniquely increases CAR T-cell functionality (**Fig. 2**) without compromising cell viability (**Fig. 3**) in both CD28- and 4-1BB-costimulated CAR T cells, our data demonstrate that low-strength IFN-I signaling promotes CAR T-cell efficacy *in vitro* by balancing function with persistence. Based on these *in vitro* conclusions, we hypothesized that the IFNα-lo condition uniquely optimizes CAR T-cell treatment efficacy *in vivo*.

To test our hypothesis, we analyzed the efficacy of CAR T cells prepared from the three culturing conditions in a xenograft lymphoma mouse model based on the OCI-Ly8 cell line (**Fig. 4A**). To control for donor-donor heterogeneity, CAR T cells were derived from multiple independent donors (three for CD28-costimulated CAR T cells and two for 4-1BB-costimulated CAR T cells). Across donors, CAR T-cell injection prolonged survival relative to the PBS-only control. For CD28-costimulated CAR T cells with donors 1 and 2, CAR T cells from the IFNα-lo condition significantly prolonged survival and slowed down tumor progression relative to CAR T cells from the IFNα-neg and IFNα-hi conditions (**Fig. 4B** and **C**, Supplementary Fig. S7A). For donor 3, although the differences between CAR T cells from IFNα-lo and IFNα-hi conditions were relatively modest compared to other donors, one mouse treated with CAR T cells from the IFNα-lo condition showed a significantly better and sustained response (Supplementary Fig. S7B and C). For 4-1BB-costimulated CAR T cells with donors 4 and 5, CAR T cells from the IFNα-lo condition consistently yielded superior outcomes compared to CAR T cells from the IFNα-neg and IFNα-hi conditions, leading to improved survival and tumor control (**Fig. 4C** and **D**, Supplementary Fig. S7D). Taken together, our findings support our hypothesis and indicate that CAR T cells generated from the IFNα-lo condition, regardless of costimulatory domain, exhibit enhanced *in vivo* efficacy compared to those from IFNα-neg or IFNα-hi conditions across multiple donors.

**Figure 4.**
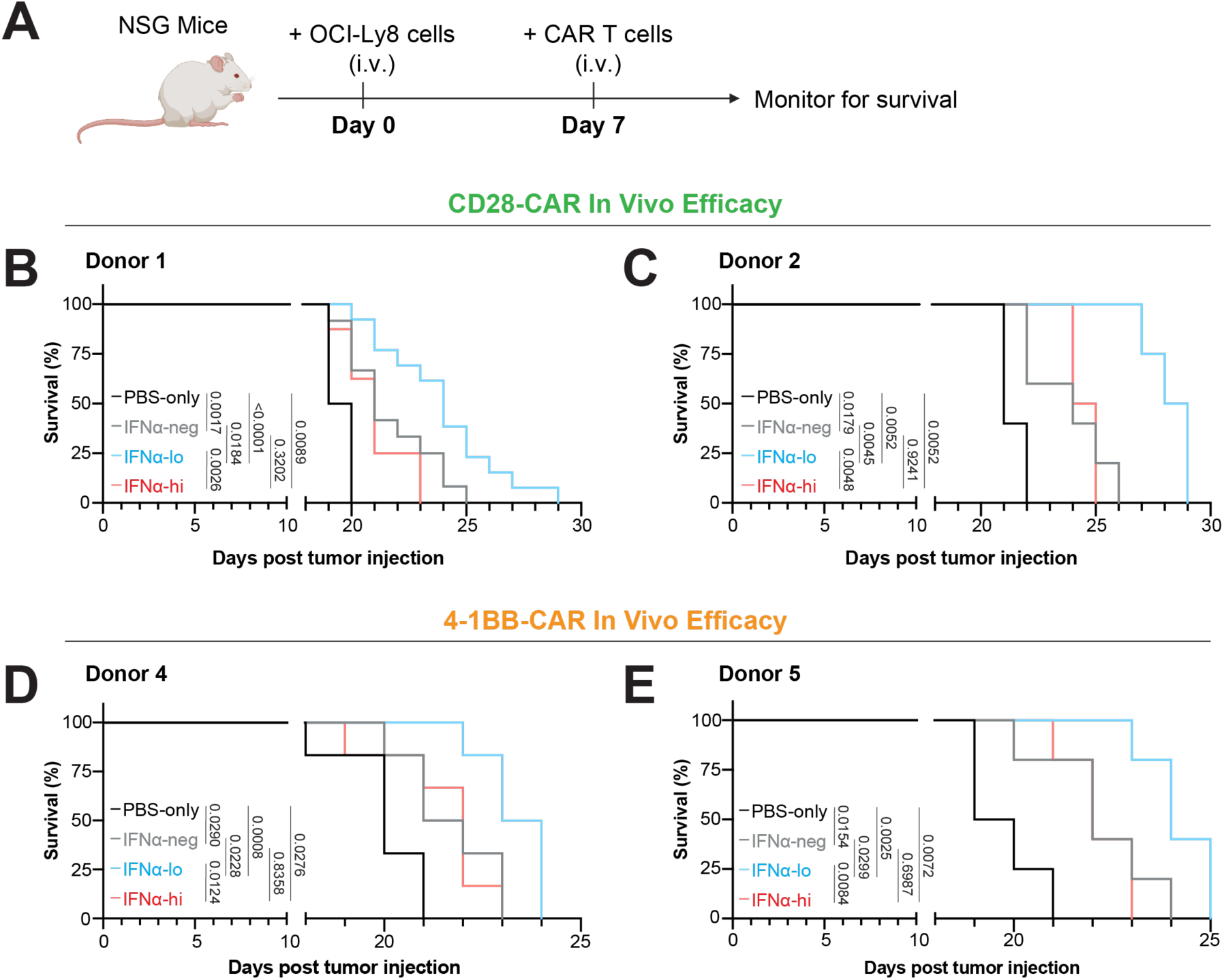
Low-strength IFN-I signaling promotes CAR T-cell treatment efficacy *in vivo*. **A,** Schematic depicting timeline for NSG mouse model based on OCI-Ly8 lymphoma cell line. Seven days following OCI-Ly8 injection, mice were randomly divided into groups that were treated with either PBS or CAR T cells from the three culturing conditions. **B** and **C,** Kaplan-Meier analysis of survival of OCI-Ly8-bearing mice treated with PBS or 5×10^5^ CD28-costimulated CAR T cells from various culture conditions. CAR T cells were generated from donor 1 (**B**, n=8,12,13,8), and donor 2 (**C**, n=5,5,4,4). **D** and **E,** Kaplan-Meier analysis of survival of OCI-Ly8-bearing mice treated with PBS or 5×10^5^ (**D**) or 1×10^6^ (**E**) 4-1BB-costimulated CAR T cells from various culture conditions. CAR T cells were generated from donor 4 (**D**, n=6,6,6,6), and donor 5 (**E**, n=4,5,5,5). Statistical analysis were performed by log-rank (Mantel-Cox) test.

### Low-strength IFN-I signaling promotes CAR T-cell efficacy in other CD19^+^ tumor models

We hypothesized low-strength IFN-I signaling can also enhance CAR T-cell efficacy against other CD19⁺ tumors. To test this, we evaluated the cytotoxicity of CAR T cells generated under different IFN-α2 treatment conditions against two additional CD19⁺ tumor cell lines: VAL (a DLBCL cell line) and NALM6 (a leukemia cell line). Consistently, IFN-I treatment enhanced CAR T-cell cytotoxicity for both CD28- and 4-1BB-costimulated CAR constructs in a dose-dependent manner (**Fig. 5A** and **D**, Supplementary Fig. S8A and D). Moreover, low-strength IFN-I signaling did not increase apoptosis or compromise CAR T-cell persistence, whereas high-strength IFN-I signaling did, as assessed *in vitro* (**Fig. 5B, C, E**, and **F**, Supplementary Fig. S8B, C, E, and F).

**Figure 5.**
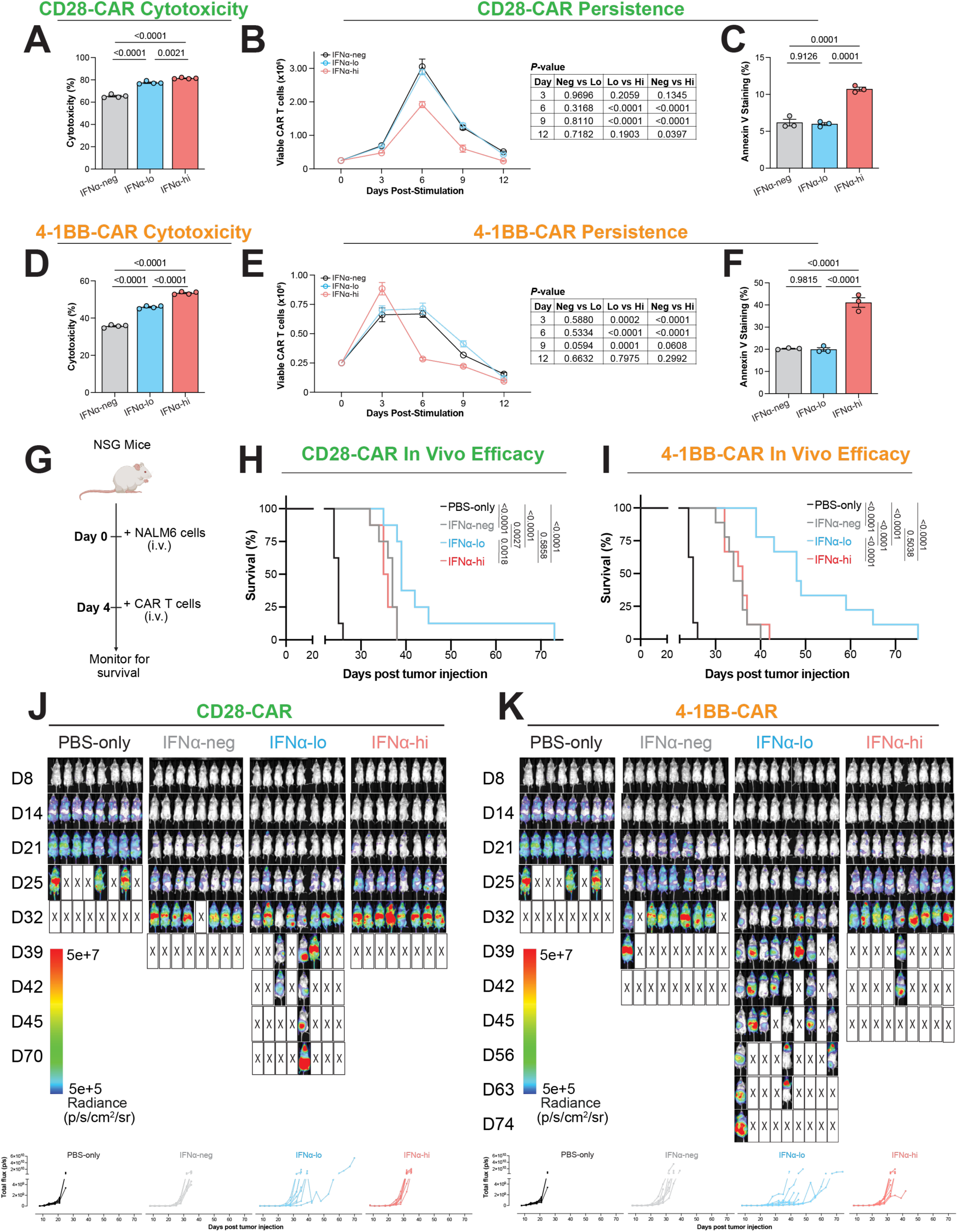
Low-strength IFN-I signaling promotes CAR T-cell efficacy in a leukemia model. **A** and **D,** Direct cytotoxicity of CD28-costimulated (**A**, n=3) and 4-1BB-costimulated (**D**, n=3) CAR T cells against firefly luciferase-expressing NALM6 target cells at E/T ratio of 1:1 via 6-hr bioluminescence assay. **B** and **E,** Line graphs depicting viable CD28-costimulated (**B**, n=3) and 4-1BB-costimulated (**E**, n=3) CAR T-cell cellularity *in vitro* following coculture with NALM6 at E/T ratio of 1:5. **C** and **F,** Apoptosis of CD28-costimulated (**C**, n=3) and 4-1BB-costimulated (**F**, n=3) CAR T cells from the three culturing conditions at the time points when cellularity declined in (**B**) and (**E**). Day 6 and day 3 after coculture, respectively. **G,** Schematic depicting timeline for NSG mouse model based on NALM6 leukemia cell line. **H** and **I,** Kaplan-Meier analysis of survival of NALM6-bearing mice treated with PBS, 5×10^5^ CD28-costimulated CAR T cells, or 1×10^6^ 4-1BB-costimulated CAR T cells, generated from a healthy donor under the three IFNα culturing conditions. **H**, mice treated with PBS or 5×10^5^ CD28-costimulated CAR T cells (n=8,8,8,8). **I,** mice treated with PBS or 1×10^6^ 4-1BB-costimulated CAR T cells (n=8,9,9,9). The PBS-treated mice in **H** and **I** were the same cohort and were used as a shared control in the same experiment. **J** and **K,** Representative bioluminescence imaging of tumor burden in mice shown in **H** (**J**) and **I** (**K**) (top), with corresponding quantification of bioluminescence (bottom). Statistical analysis were performed via one-way (**A**, **C**, **D**, and **F**) or two-way (**B** and **E**) ANOVA with Tukey’s correction for multiple comparisons, or log-rank (Mantel-Cox) test (**H** and **I**). **A**-**F**, All data are shown as mean ± SEM of n experimental replicates from a representative donor. Results are representative of independent experiments using CAR T cells from 3 different donors.

Furthermore, in an NALM6-based *in vivo* xenograft leukemia model, both CD28- and 4-1BB-costimulated CAR T cells generated under IFNα-lo conditions outperformed the other conditions, as evidenced by prolonged survival and reduced tumor burden (**Fig. 5G-K**). Together, these data suggest that the IFNα-lo approach has broad applicability for enhancing CAR T-cell efficacy against other CD19⁺ tumors.

### Low-strength IFN-I signaling promotes durable CAR T-cell efficacy

To rigorously evaluate the durability of antitumor responses mediated by CAR T cells generated under IFNα-lo conditions, we refined the OCI-Ly8 xenograft model by introducing a secondary tumor challenge following initial tumor control with CAR T-cell therapy. This approach enabled assessment of the sustained therapeutic efficacy of these CAR T cells in the setting of residual or recurrent disease (**Fig. 6A**). CAR T cells generated under IFNα-lo conditions conferred superior survival benefit and tumor control compared with those generated under IFNα-neg conditions (**Fig. 6B-D**). This was accompanied by higher levels of CAR T cells in the peripheral blood and spleen at later time points following the second challenge (**Fig. 6E** and **F**), although memory phenotypes were not markedly altered (Supplementary Fig. S9). Collectively, these data suggest that the IFNα-lo approach promotes durable CAR T-cell efficacy.

**Figure 6.**
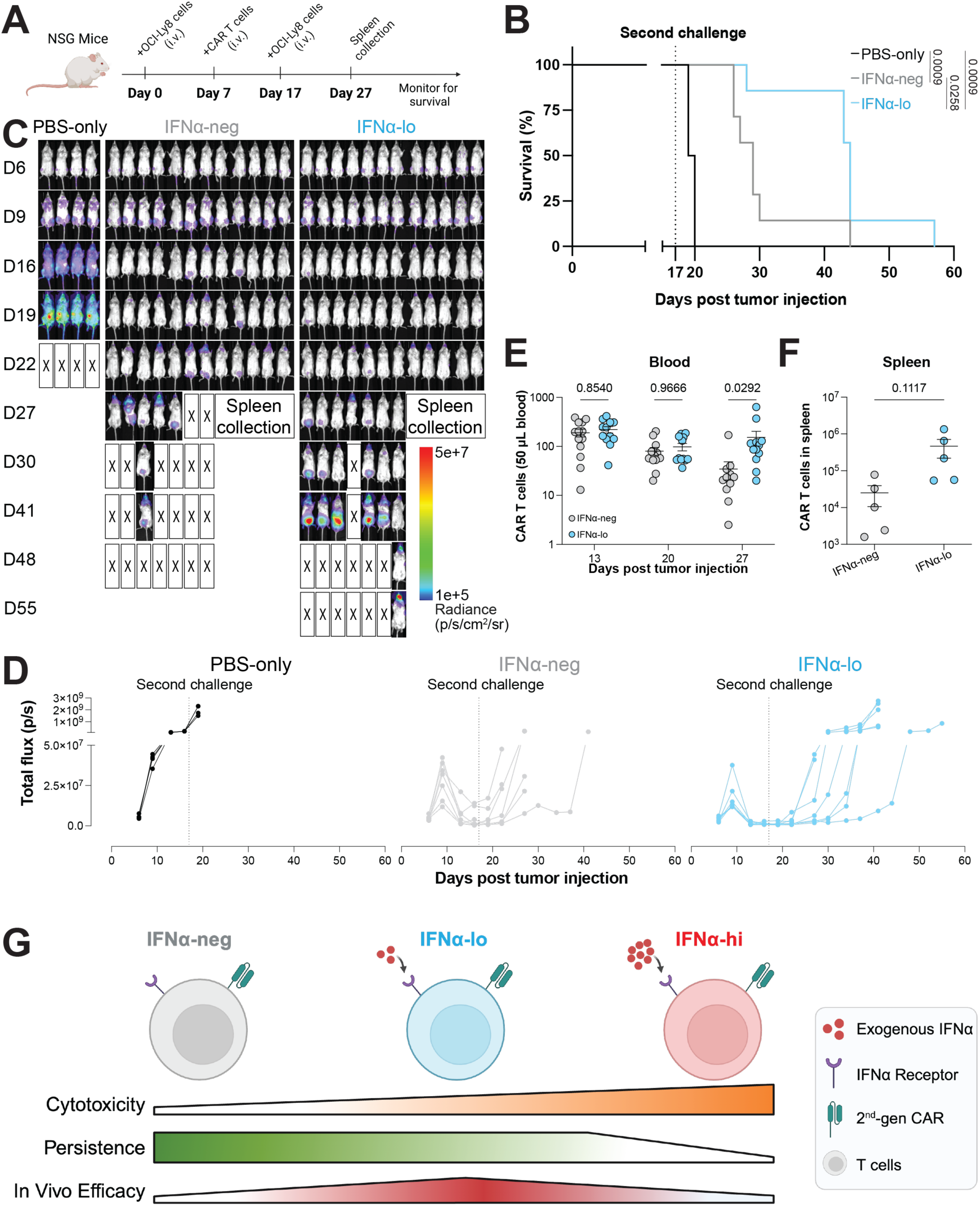
Low-strength IFN-I signaling promotes durable CAR T-cell efficacy. **A,** Schematic depicting timeline for OCI-Ly8-based xenograft tumor model used to assess durable CAR T-cell efficacy. **B,** Kaplan-Meier analysis of survival of OCI-Ly8-bearing mice treated with PBS or 3×10^6^ CD28-costimulated CAR T cells generated from a healthy donor under 2 IFNα culturing conditions (n=4,7,7). **C** and **D,** Representative bioluminescence imaging of tumor burden in mice shown in **B**, along with additional CAR T-cell-treated mice used for spleen collection (**C**, n=4,12,12), and corresponding quantification of bioluminescence signals for mice in (**B**, bottom, n=4,7,7). **E,** Longitudinal quantification of CAR^+^ cells in 50 μL peripheral blood from mice in **C**. **F,** Quantification of CAR^+^ cells in the spleen collected at day 27 from mice in **C**. **G,** Cartoon summarizing effects of IFN-I signaling on CAR T cells at different doses. Statistical analysis were performed by log-rank (Mantel-Cox) test (**B**), two-way (**E**) ANOVA with Tukey’s correction for multiple comparisons, or unpaired t-test (**F**).

In conclusion, our *in vitro* and *in vivo* studies support a model in which low-strength IFN-I signaling selectively promotes CAR T-cell treatment efficacy (**Fig. 6G**). IFN-I signaling promotes CAR T-cell functionality in a concentration-dependent manner, leading to more effective cytotoxicity, cytokine release, and tumor control. However, at a high enough concentration, IFN-I signaling also increases T-cell apoptosis, leading to reduced yield, poorer persistence, and ineffective tumor control. Although the optimal level of IFN-I signaling may be subject to donor-donor heterogeneity, CAR T cells treated with low-strength IFN-I signaling, regardless of costimulatory domain, consistently exhibit enhanced effector functions against lymphoma and leukemia with durable responses.

## DISCUSSION

In this study, we leveraged single-cell transcriptomic analyses of patient IPs from two independent datasets, *in vitro* functional and phenotypic analyses, and *in vivo* xenograft mouse studies to establish that low-strength IFN-I signaling during the *ex vivo* CAR T-cell manufacturing process selectively promotes CAR T-cell treatment efficacy *in vivo* in a costimulation-independent manner. Our strategy of harnessing IFN-I signaling to enhance CAR T-cell efficacy is grounded in clinical data and existing literature, while also highlighting IFN-I as an enhancer with significant translational potential for use during the CAR T-cell manufacturing process.

There are major attractive qualities that make IFN-I a uniquely promising and translatable enhancer for both CAR T-cell therapy and other emerging cellular therapy modalities. Unlike all other natural and engineered cytokines being developed as CAR T-cell enhancers (including IL-7, IL-10, IL-15, IL-21, Fc-IL-4)^39–43^, IFN-Is are uniquely biocompatible drugs with an existing and extensive clinical track record for the treatment of malignancies^22^, autoimmune diseases^23^, and viral infections^24^. First discovered in the 1950s for their antiviral activities, IFN-I is one of only two cytokines ever approved by the FDA for cancer treatment (the other being IL-2).^44^ Biologically, IFN-I elicits a known signaling pathway through the type I interferon receptor to induce antiviral interferon-stimulated genes.^13,15^ Their pharmacodynamics and pharmacokinetics have been thoroughly explored in pre-clinical and clinical studies.^45^ Importantly, whereas existing cytokines such as IL-15 and IL-21 boost stem cell memory phenotype^39^ and persistence^41^, respectively, our IFN-I conditioning strategy boosts immediate effector functions, including cytotoxicity and cytokine production. Thus, IFN-I enables a distinct and complementary enhancement strategy that synergizes with the current state of the art. Our novel manufacturing method was mechanistically inspired by bioinformatics findings from clinical biospecimens, allowing it to recapitulate a physiological and clinically relevant phenotype. By utilizing a biologically, pharmacologically, and clinically characterized pharmacophore, our low-strength IFN-I manufacturing approach is both straightforward and safe. Our approach involves adding IFN-I cytokines during the manufacturing process, followed by IFN-I removal before CAR T-cell infusion for treatment. This design fundamentally differs from sustained IFN-I exposure^18^, allowing us to avoid T-cell exhaustion and harness the beneficial effects of low-strength IFN-I signaling during *ex vivo* CAR T-cell manufacture. Importantly, restricting IFN-I exposure to the manufacturing process ensures simplicity and compatibility with existing manufacturing methods and CAR constructs, while eliminating potential *in vivo* IFN-I-associated toxicities.

In terms of basic biology, we found that IFN-I signaling enhances CAR T-cell cytotoxicity in a dose-dependent manner; however, excessively strong IFN-I signaling also increases apoptosis, thereby reducing persistence. Our findings are compatible with existing literature on T-cell-intrinsic effects of IFN-I signaling. Enhancement of cytotoxicity and type I cytokine secretion among CD8^+^ and CD4^+^ T cells after culturing with exogenous IFN-I is well-established.^16,46^ Moreover, the field’s current understanding is that T-cell-intrinsic IFN-I signaling leads to different consequences, depending on whether TCR stimulation temporally precedes (“in-sequence”) or follows (“out-of-sequence”) IFN-I signaling.^13,47^ In-sequence signaling generally enhances T-cell functions, while out-of-sequence signaling generally does the opposite.^13^ In our studies, since T-cell activation (via CD3/CD28 Dynabeads) preceded IFN-I treatment, our CAR T cells were subject to in-sequence signaling under the current paradigm. Moreover, our findings mirror the expected consequences of in-sequence IFN-I signaling. Lastly, the concentration-dependent nature of our model is compatible with other studies demonstrating that different cellular functions exhibit varying sensitivities to IFN-I signaling.^33^

Our findings help reconcile previously conflicting reports regarding the role of IFN-I in CAR T-cell function. While some studies report that IFN-I signaling impairs CAR T-cell efficacy^20,21,48^, others report beneficial effects^49^. Jung et al.^48^ and Evgin et al.^21^ report that IFN-I signaling impairs CAR T-cell efficacy. Both studies used IFN-β, which has up to 30-fold higher affinity for the type I interferon receptor compared to IFN-α2.^50^ Considering the high-affinity IFN-I subtype and concentrations applied, their experimental conditions align with our IFNα-hi condition, providing a plausible explanation for their detrimental outcomes. Similarly, Harrer et al. report that autocrine IFN-I signaling (through CAR-activation-induced secretion of IFN-α2 or IFN-β) impairs CAR T-cell efficacy.^20^ The resulting IFN-I bioactivity in their cell supernatant aligns with our IFNα-hi condition, triggering increased CAR T-cell apoptosis. In contrast, Zhao et al. reported that IFN-I signaling enhances CAR T-cell efficacy.^49^ We reason that their beneficial findings reflect a weak endogenous IFN-I stimulus, as no exogenous IFN-I was added in their *in vitro* system. Together, these comparisons highlight how our model, which defines a non-linear relationship between IFN-I signaling strength and CAR T-cell efficacy, provides a unifying framework to reconcile previously conflicting reports.

Our study advances the understanding of IFN-I signaling in T cells and lays the foundation for translational applications in CAR T-cell therapies, though several avenues remain for exploration. Since IFN-I signaling enhances CAR T-cell cytokine production (IL-2, TNF, IFN-γ), it may theoretically increase the risk of cytokine release syndrome (CRS) and immune effector cell-associated neurotoxicity syndrome (ICANS).^51^ The xenograft mouse model is not suitable for evaluating either toxicities.^52^ Any future application of our IFN-I strategy in the clinical trials context should carefully titrate CAR T-cell dosages, consider patient performance status, monitor treatment toxicities, implement evidence-based clinical management (e.g., tocilizumab for CRS, dexamethasone for ICANS), and incorporate rapid reversal mechanisms where appropriate (e.g., suicide switches for CAR T cells). Furthermore, we tested only two IFN-I concentrations, emphasizing the need for more rigorous dosing studies to determine the optimal therapeutic window. Finally, although our study focused on IFN-α2 due to its clinical relevance, it underscores the broader potential of the IFN-I family, with many subtypes still underexplored and potentially capable of further enhancing CAR T-cell functionality through cytokine modulation.

Looking ahead, the simplicity, clinical familiarity, and GMP compatibility of low-strength IFN-I signaling make this strategy readily adoptable in clinical CAR T-cell manufacturing. Although our studies center on CD19-directed CAR T cells and B cell malignancies, our approach is likely generalizable across additional CAR targets and disease settings. These findings provide strong rationale for advancing low-strength IFN-I signaling toward clinical trials evaluation as a scalable and broadly applicable manufacturing strategy to enhance cell therapies.

## ACKNOWLEDGEMENTS

We thank the NIH New Innovator award (1DP2AI144245), NIH R21AI169159, the American Cancer Society Scholar Award (SG-22-136-01-1BCD), and NIH R61AG090388 for financial support of this research, including support for reagents and personnel (to J.H.). Y.H. was supported by the University of Chicago MSTP Training Grant (T32GM007281). N.W.A. was supported by the University of Chicago MTCR Training Grant (T32CA009594). Flow cytometry was performed at the Cytometry and Antibody Technology Facility at the University of Chicago, which receives financial support from the Cancer Center Support Grant (P30CA014599). Live bioluminescence imaging was performed at the University of Chicago Integrated Small Animal Imaging Research Resource. We thank the UChicago Blood Donation Center for healthy donor PBMCs. We thank Aniruddhsingh Solanki at the Animal Resource Center at University of Chicago for excellent technical assistance. We thank Dr. Alexandra Rojek from Dr. Justin Kline’s lab for sharing the VAL cell line. We also acknowledge Ali Rahman for assistance with cell culture, molecular cloning, and the optimization of immunoassay conditions.

## AUTHOR CONTRIBUTIONS

**E. Tang**: Conceptualization, data curation, formal analysis, investigation, validation, methodology, bioinformatics analysis, writing-original draft, writing-review and editing. **Y. Hu**: Conceptualization (original project idea), clinical data collection, data curation, formal analysis, investigation, validation, methodology, writing-original draft, writing-review and editing. **G. Cao**: Conceptualization, investigation, bioinformatics analysis. **D. Nguyen**, **N. W. Asby**, **N. S. Aboelella**, and **H. A. Ruiz**: Investigation and methodology. **X. Cai**, **W. Zhang**, **Y. Zhao**, **L. Xie**, and **X. Chen**: Methodology. **M. R. Bishop**, **P. A. Riedell**, **J. L. LaBelle**, and **J. P. Kline**: Methodology and resources. **J. Huang**: Conceptualization, resources, funding acquisition, supervision, formal analysis, methodology, writing-original draft, writing-review and editing, project administration.

## Notes

### Competing Interest Statement

J.H., Y.H., and E.T. are listed as inventors on a patent related to interferon-enhanced CAR-T cells (US provisional patent application 63/789,043). M.R.B. reports membership on an Advisory Board or Consultancy for Kite/Gilead, Novartis, CRISPR Therapeutics, Autolus Therapeutics, BMS, Incyte, Sana Biotechnology, Iovance Biotherapeutics. He has served on a Speakers Bureau for BMS, Kite/Gilead, Agios, and Incyte. P.A.R. reports Research Support/Funding: BMS, Kite Pharma, Inc./Gilead, MorphoSys, Calibr, Tessa Therapeutics, Fate Therapeutics, Xencor, and Novartis Pharmaceuticals Corporation. Speakers Bureau: Kite Pharma, Inc./Gilead; Consultancy on advisory boards: AbbVie, Novartis Pharmaceuticals Corporation, BMS, Janssen, BeiGene, Karyopharm Therapeutics Inc., Takeda Pharmaceutical Company, Kite Pharma, Inc./Gilead, Sana Biotechnology, Nektar Therapeutics, Nurix Therapeutics, Intellia Therapeutics, and Bayer. Honoraria: Novartis Pharmaceuticals Corporation. J.L.L. reports other grants from AbbVie and the American Cancer Society outside the submitted work. J.P.K. receives research support from Merck, Verastem, and iTeos; has served on a speakers bureau for Kite/Gilead; and has served on advisory boards for Verastem, Seattle Genetics, MorphoSys, and Karyopharm. No disclosures were reported by the other authors.

### Summary of Updates

Two more figures have been included.

