## Supplemental Data for "Low-Strength Type I Interferon Signaling Promotes CAR T-Cell Treatment Efficacy"

### RUNNING TITLE

Low-Strength IFN-I Enhances CAR T Cells

### DECLARATIONS OF INTERESTS

J.H., Y.H., and E.T. are listed as inventors on a patent related to interferon-enhanced CAR-T cells (US provisional patent application 63/789,043). M.R.B. reports membership on an Advisory Board or Consultancy for Kite/Gilead, Novartis, CRISPR Therapeutics, Autolus Therapeutics, BMS, Incyte, Sana Biotechnology, Iovance Biotherapeutics. He has served on a Speakers Bureau for BMS, Kite/Gilead, Agios, and Incyte. P.A.R. reports Research Support/Funding: BMS, Kite Pharma, Inc./Gilead, MorphoSys, Calibr, Tessa Therapeutics, Fate Therapeutics, Xencor, and Novartis Pharmaceuticals Corporation. Speakers Bureau: Kite Pharma, Inc./Gilead; Consultancy on advisory boards: AbbVie, Novartis Pharmaceuticals Corporation, BMS, Janssen, BeiGene, Karyopharm Therapeutics Inc., Takeda Pharmaceutical Company, Kite Pharma, Inc./Gilead, Sana Biotechnology, Nektar Therapeutics, Nurix Therapeutics, Intellia Therapeutics, and Bayer. Honoraria: Novartis Pharmaceuticals Corporation. J.L.L. reports other grants from AbbVie and the American Cancer Society outside the submitted work. J.P.K. receives research support from Merck, Verastem, and iTeos; has served on a speakers bureau for Kite/Gilead; and has served on advisory boards for Verastem, Seattle Genetics, MorphoSys, and Karyopharm. No disclosures were reported by the other authors.

### SUPPLEMENTARY TABLES AND FIGURES

#### Table S1. Patient characteristics and relevant clinical data.

| **Characteristics** | **CR-1** | **CR-2** | **CR-3** | **CR-4** | **CR-5** | **N-CR-1** | **N-CR-2** | **N-CR-3** |
|---|---|---|---|---|---|---|---|---|
| Age Decade, y | 50s | 70s | 50s | 40s | 60s | 70s | 60s | 50s |
| Sex | Female | Male | Female | Male | Male | Male | Female | Male |
| Disease stage | I | II | II | IV | IV | IV | III | I |
| No. of prior therapies | 4 | 2 | 2 | 5 | 5 | 3 | 3 | 3 |
| Prior lines of therapy | Radiation | R-CHOP | DA-EPOCH w/ IT MTX | R-CHOP | R-CHOP | BR | BR | R-CHOP |
| | R-CHOP | R-ICE | ICE | R-ICE | Benda-obinutuzumab | DA-R-EPOCH | ICE | DA-R-EPOCH |
| | R-ICE | | | BEAM + ASCT | R-ICE | Hu5F9-G4 + rituximab | GEMOX | R-ICE |
| | BEAM + ASCT | | | FCR + AlloSCT | Hu5F9-G4 + rituximab | | | |
| | | | | R-GEMOX | R-GEMOX | | | |
| Disease status | Relapsed | Relapsed | Primary refractory | Relapsed | Relapsed, then refractory | Primary refractory | Relapsed, then refractory | Primary refractory |
| **Baseline (Ref. range)** | | | | | | | | |
| LDH, U/L (116-245 U/L) | 186 | 165 | 312 | 203 | 311 | 332 | 336 | 260 |
| CRP, mg/dL (<0.5 mg/dL) | 0.6 | 0 | 3.5 | 0.3 | ND | 0.6 | 0 | 0.2 |
| Ferritin, ng/mL (20-300 ng/mL) | 215 | 110 | 576 | 544 | ND | 64 | 124 | 294 |
| ECOG PS | 0 | 1 | 2 | 1 | 1 | 0 | 1 | 1 |
| Bridging therapy | No | No | Yes | Yes | No | No | Yes | No |
| Maximum Grade CRS | 1 | 2 | 2 | 1 | 1 | 1 | 1 | 0 |
| Maximum Grade ICANS | 4 | 2 | 3 | 1 | 3 | 0 | 0 | 0 |
| Tocilizumab or steroids | Steroids | Both | Both | Tocilizumab | Both | Neither | Tocilizumab | Neither |

Abbreviations: ASCT, autologous stem cell transplant; AlloSCT, allogeneic stem cell transplant; BEAM, BCNU (carmustine), etoposide, cytarabine, melphalan; BR, bendamustine, rituximab; CRP, C-reactive protein; CRS, cytokine release syndrome; DA-EPOCH, dose-adjusted etoposide, prednisone, oncovin, cyclophosphamide, hydroxydaunorubicin; DA-R-EPOCH, dose-adjusted rituximab, etoposide, prednisone, oncovin, cyclophosphamide, hydroxydaunorubicin; ECOG PS, Eastern Cooperative Oncology Group Performance Score; FCR, fludarabine, cyclophosphamide, rituximab; GEMOX, gemcitabine, oxaliplatin; Hu5F9-G4, magrolimab; ICANS, immune effector cell-associated neurotoxicity syndrome; ICE, ifosfamide, carboplatin, etoposide; IT MTX, intrathecal methotrexate; LDH, lactate dehydrogenase; ND, not determined; R-CHOP, rituximab, cyclophosphamide, hydroxydaunorubicin, oncovin, prednisone; R-GEMOX, rituximab, gemcitabine, oxaliplatin; R-ICE, rituximab, ifosfamide, carboplatin, etoposide

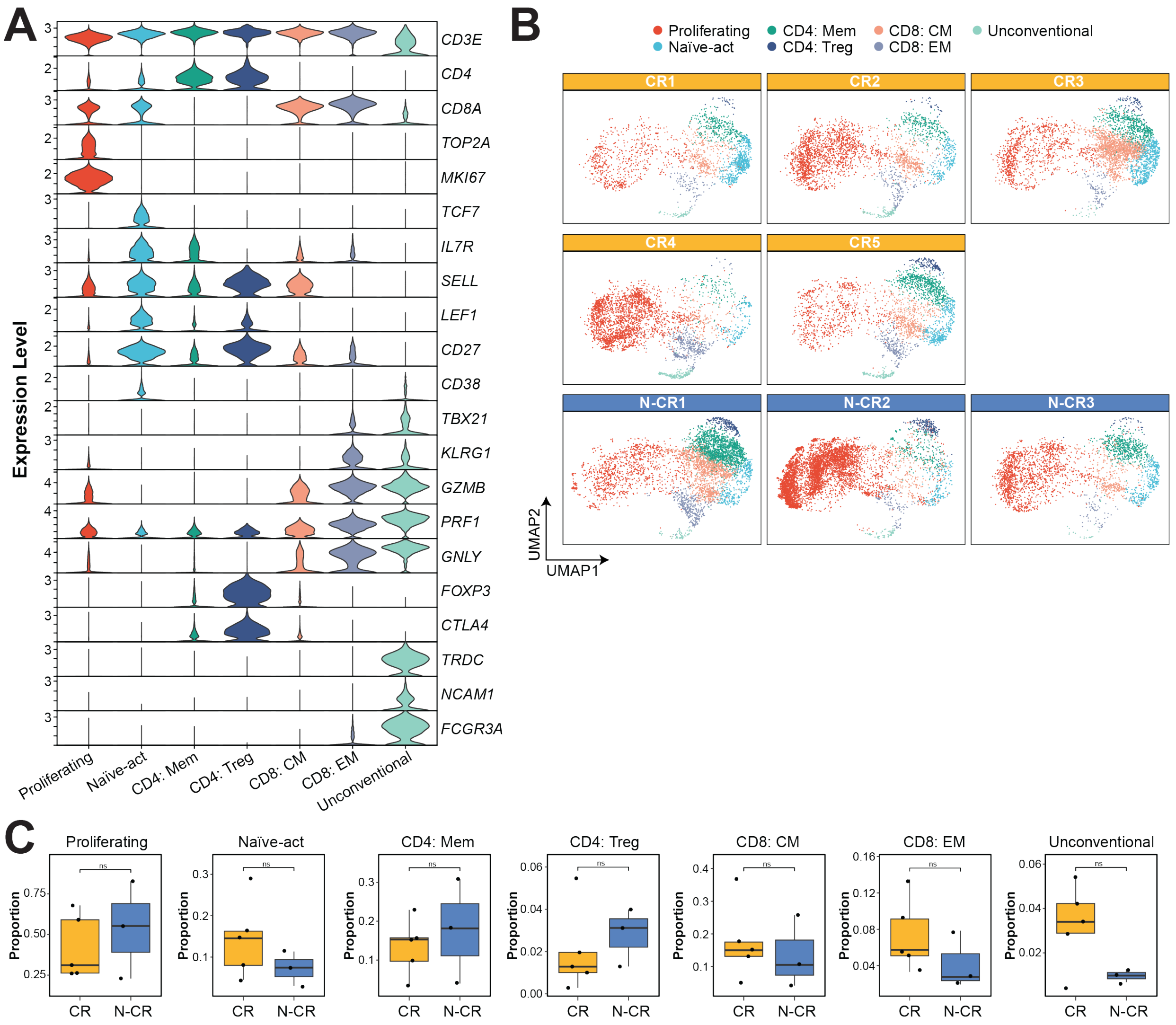


**Figure S1. Transcriptomic analysis of T cells in discovery dataset.**
**A,** Violin plots depicting normalized expression levels of markers for each cluster. **B,** UMAPs demonstrating the distributions of the 7 T-cell clusters in each patient. **C,** Box plots depicting proportions of each phenotype within CR and N-CR T cells. Each dot represents a measurement from a single patient. Proportions in CR and N-CR T cells were compared by Wilcoxon rank-sum test, whereby ns indicates no significance.

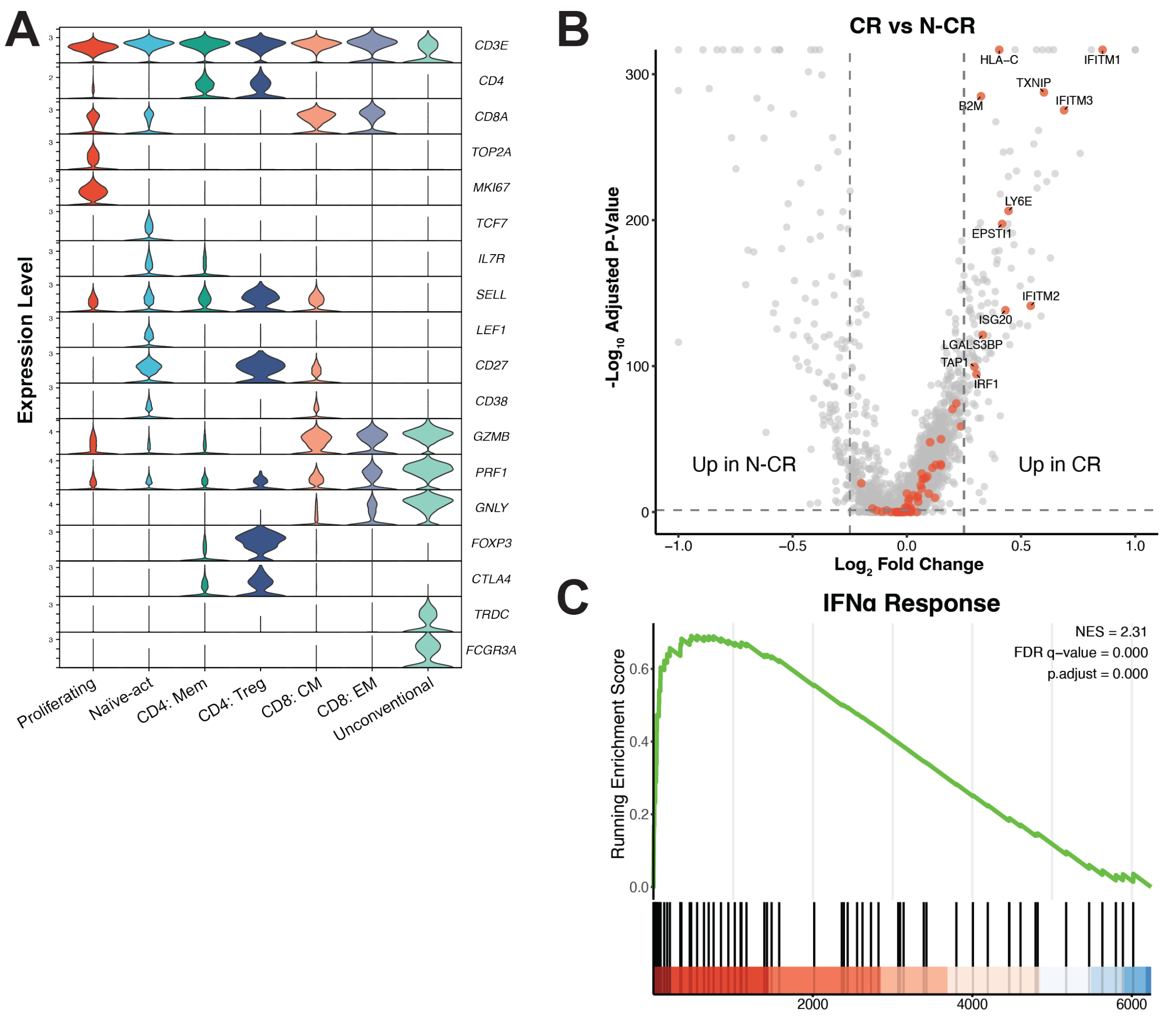


**Figure S2. Transcriptomic analysis of T cell in validation dataset.**

**A,** Violin plots depicting normalized expression levels of markers for each cluster. **B,** Volcano plot depicting DEGs between CR and N-CR across T cells. Genes involved in IFNα response gene set are colored in red. Dotted lines represent cutoff values for DEGs ($log_2$ fold-change of ± 0.25) and significance threshold (adjusted p-value of 0.05). **C,** Enrichment plots for the IFNα response gene set differentially expressed between CR and N-CR T cells.

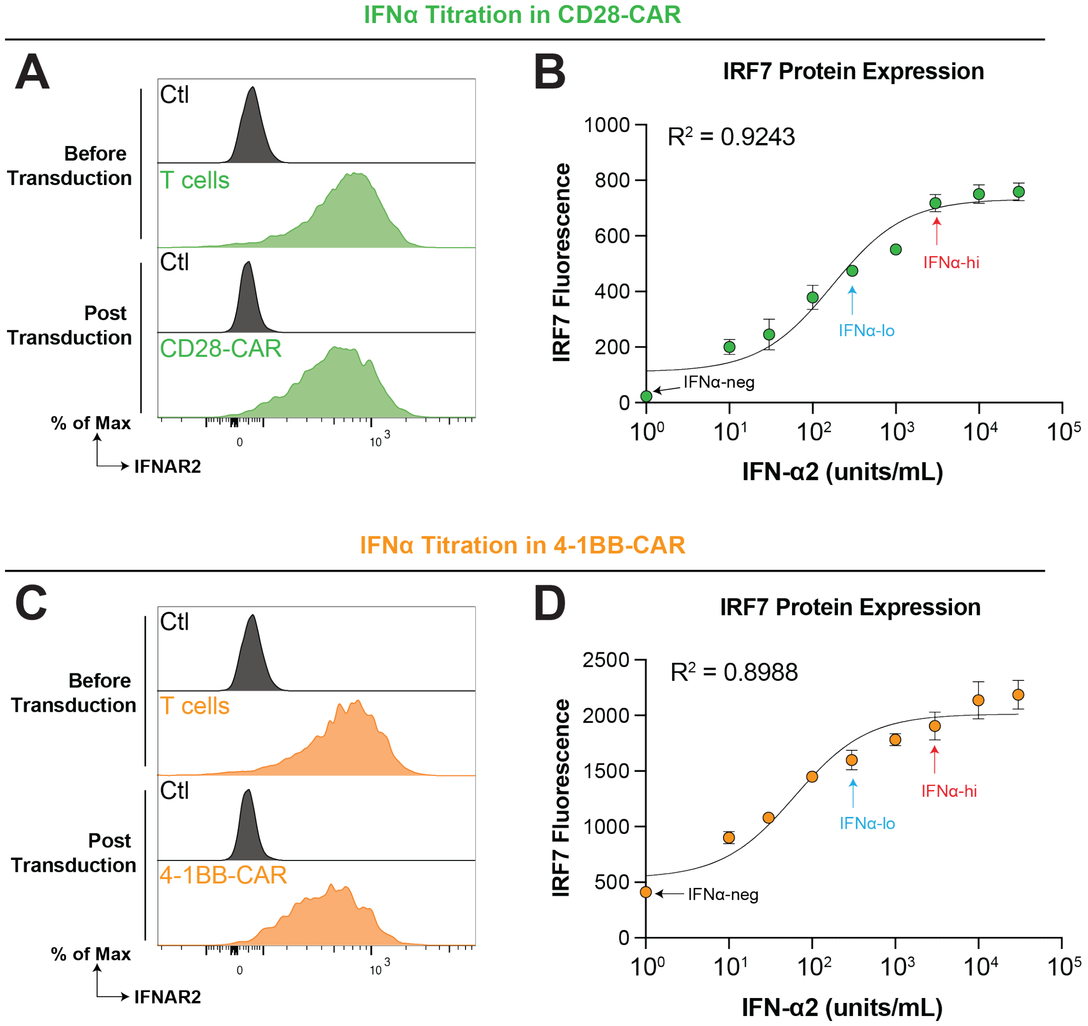


**Figure S3. Type I interferon receptor expression and IFN-α2 titration in CAR T cells.**
**A** and **C,** Representative flow plots showing expression of type I interferon receptor (IFNAR2) on T cells before (top) or post (bottom) CAR transduction. T cells were transduced with CD28-costimulated (**A**) or 4-1BB-costimulated (**C**) CARs. An isotype control antibody was used as the negative control for $IFNAR2^+$ gating. **B** and **D,** Titration of IFN-α2 to CD28-costimulated (**B,** n=3) and 4-1BB-costimulated (**D,** n=3) CAR T cells. Cells were subjected to IFN-α2 for 18 hours prior to intracellular staining for IRF7, a known ISG. The data points were fitted to a dose-response curve. The three IFN-I signaling conditions were labeled on the curve. Results are representative of independent experiments using CAR T cells from 2 different donors.

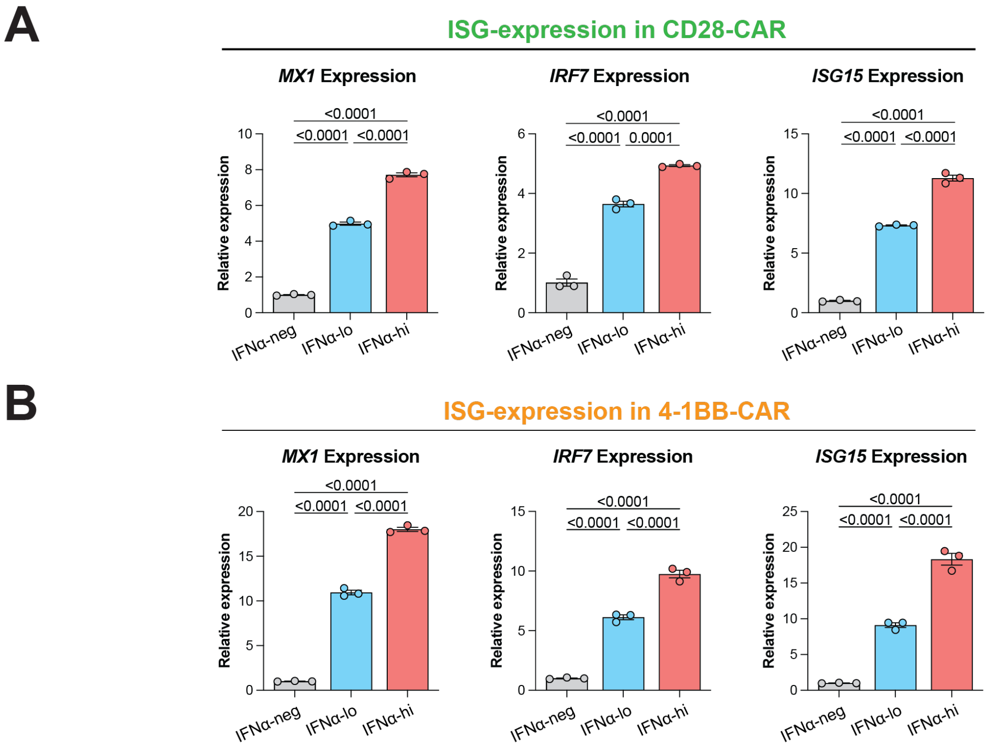


**Figure S4. Expression of ISGs with IFN-α2 signaling.**

**A** and **B,** Relative expression by quantitative reverse transcription PCR (RT-qPCR) of given ISGs in CD28-costimulated (**A,** n=3) and 4-1BB-costimulated (**B,** n=3) CAR T cells from the three culturing conditions. Expression of ISGs was normalized to the expression of β-actin transcript (*ACTB*) and subsequently to the IFNα-neg condition. All data are shown as mean ± SEM of n experimental replicates from a representative donor. Results are representative of independent experiments using CAR T cells from 2 different donors. Statistical comparisons were performed with one-way ANOVA with Tukey's correction for multiple comparisons.

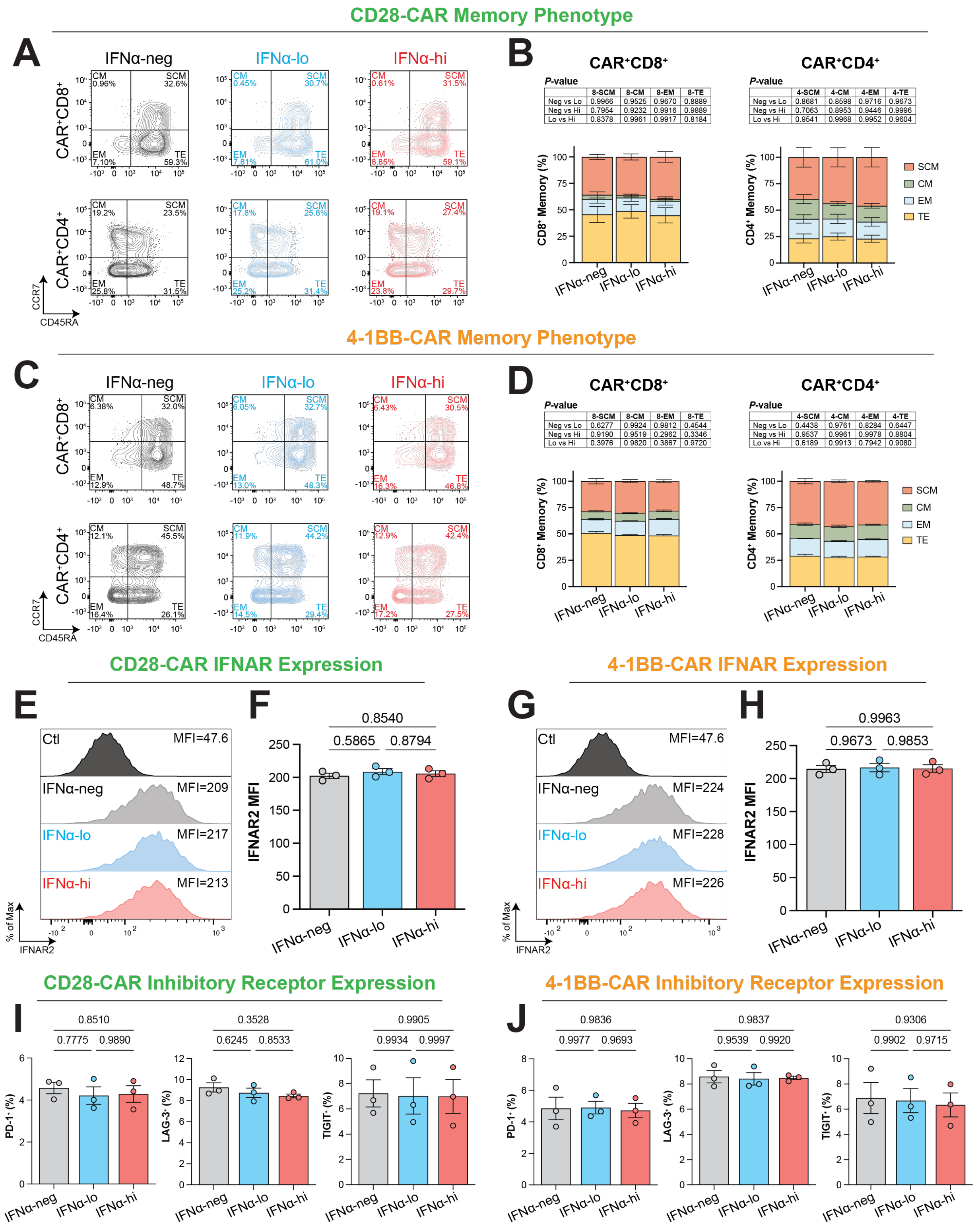
CD28-CAR Memory Phenotype
A
IFNα-neg
IFNα-lo
IFNα-hi
CAR⁺CD8⁺
CAR⁺CD4⁺
CCR7
CD45RA
B
CAR⁺CD8⁺
P-value
8-SCM 8-CM 8-EM 8-TE
Neg vs Lo 0.9966 0.9525 0.9670 0.8889
Neg vs Hi 0.7954 0.9232 0.9916 0.9889
Lo vs Hi 0.8378 0.9961 0.9917 0.8184
CAR⁺CD4⁺
P-value
4-SCM 4-CM 4-EM 4-TE
Neg vs Lo 0.8681 0.8598 0.9716 0.9673
Neg vs Hi 0.7063 0.8953 0.9446 0.9996
Lo vs Hi 0.9541 0.9968 0.9952 0.9604
CD8⁺ Memory (%)
CD4⁺ Memory (%)
SCM
CM
EM
TE
4-1BB-CAR Memory Phenotype
C
D
CAR⁺CD8⁺
P-value
8-SCM 8-CM 8-EM 8-TE
Neg vs Lo 0.6277 0.9924 0.9812 0.4544
Neg vs Hi 0.9190 0.9519 0.2962 0.3346
Lo vs Hi 0.3976 0.9820 0.3867 0.9720
CAR⁺CD4⁺
P-value
4-SCM 4-CM 4-EM 4-TE
Neg vs Lo 0.4438 0.9761 0.8284 0.6447
Neg vs Hi 0.9537 0.9961 0.9978 0.8804
Lo vs Hi 0.6189 0.9913 0.7942 0.9080
CD28-CAR IFNAR Expression
E
F
Ctl MFI=47.6
IFNα-neg MFI=209
IFNα-lo MFI=217
IFNα-hi MFI=213
% of Max
IFNAR2
IFNAR2 MFI
0.8540
0.5865
0.8794
4-1BB-CAR IFNAR Expression
G
H
Ctl MFI=47.6
IFNα-neg MFI=224
IFNα-lo MFI=228
IFNα-hi MFI=226
0.9963
0.9673
0.9853
CD28-CAR Inhibitory Receptor Expression
I
0.8510
0.7775 0.9890
0.3528
0.6245 0.8533
0.9905
0.9934 0.9997
PD-1⁺ (%)
LAG-3⁺ (%)
TIGIT⁺ (%)
4-1BB-CAR Inhibitory Receptor Expression
J
0.9836
0.9977 0.9693
0.9837
0.9539 0.9920
0.9306
0.9902 0.9715

**Figure S5. Immunophenotypic characterization of CAR T cells.**
**A** and **C,** Representative flow plots depicting gating for CD45RA and CCR7 memory markers in CD28-costimulated (**A**) and 4-1BB-costimulated (**C**) CAR T cells. Top, $CD8^+$ cells, bottom, $CD4^+$ T cells. **B** and **D,** Stacked bar plots comparing proportions of the SCM ($CD45RA^+CCR7^+$), CM ($CD45RA^-CCR7^+$), EM ($CD45RA^-CCR7^-$), and TE ($CD45RA^+CCR7^-$) memory phenotypes among $CD8^+$ (left, n=3) and $CD4^+$ (right, n=3) T cells in CD28-costimulated (**B**) and 4-1BB-costimulated (**D**) CAR T cells. SCM, stem cell memory, CM, central memory, EM, effector memory, TE, terminally differentiated effector. **E**-**H,** IFNAR2 expression in CAR T cells. **E** and **G,** Representative flow plots depicting gating for $IFNAR2^+$ cells in CD28-costimulated (**E**) and 4-1BB-costimulated (**G**) CAR T cells. **F** and **H,** Expression of IFNAR2 as measured by gMFI in CD28-costimulated (**F**) and 4-1BB-costimulated (**H**) CAR T cells from the three culturing conditions. An isotype antibody was used as the negative control for $IFNAR2^+$ gating. **I** and **J,** Bar plots showing expression of inhibitory receptors (PD-1, LAG-3, and TIGIT) in CD28-costimulated (**I**, n=3) and 4-1BB-costimulated (**J**, n=3) CAR T cells after ex vivo culturing. All data are shown as mean ± SEM of n experimental replicates from a representative donor. Results are representative of independent experiments using CAR T cells from 3 (**A**-**D**, **I**-**J**) or 2 (**E**-**H**) different donors. Statistical analysis were performed via two-way (**B** and **D**) or one-way (**F**, **H**-**J**) ANOVA with Tukey's correction for multiple comparisons.

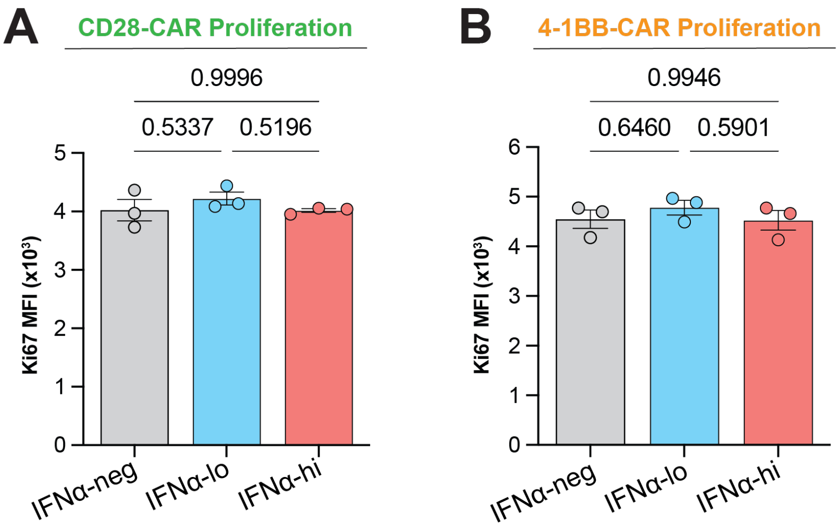


**Figure S6. CAR T-cell proliferation.**
**A** and **B,** Proliferation of CAR T cells from the three culturing conditions, as measured by Ki-67 expression (gMFI), in CD28-costimulated (**A**, n=3) and 4-1BB-costimulated (**B**, n=3) CAR T cells. All data are shown as mean ± SEM of n experimental replicates from a representative donor. Results are representative of independent experiments using CAR T cells from 2 different donors. Statistical analysis were performed via one-way ANOVA with Tukey's correction for multiple comparisons.

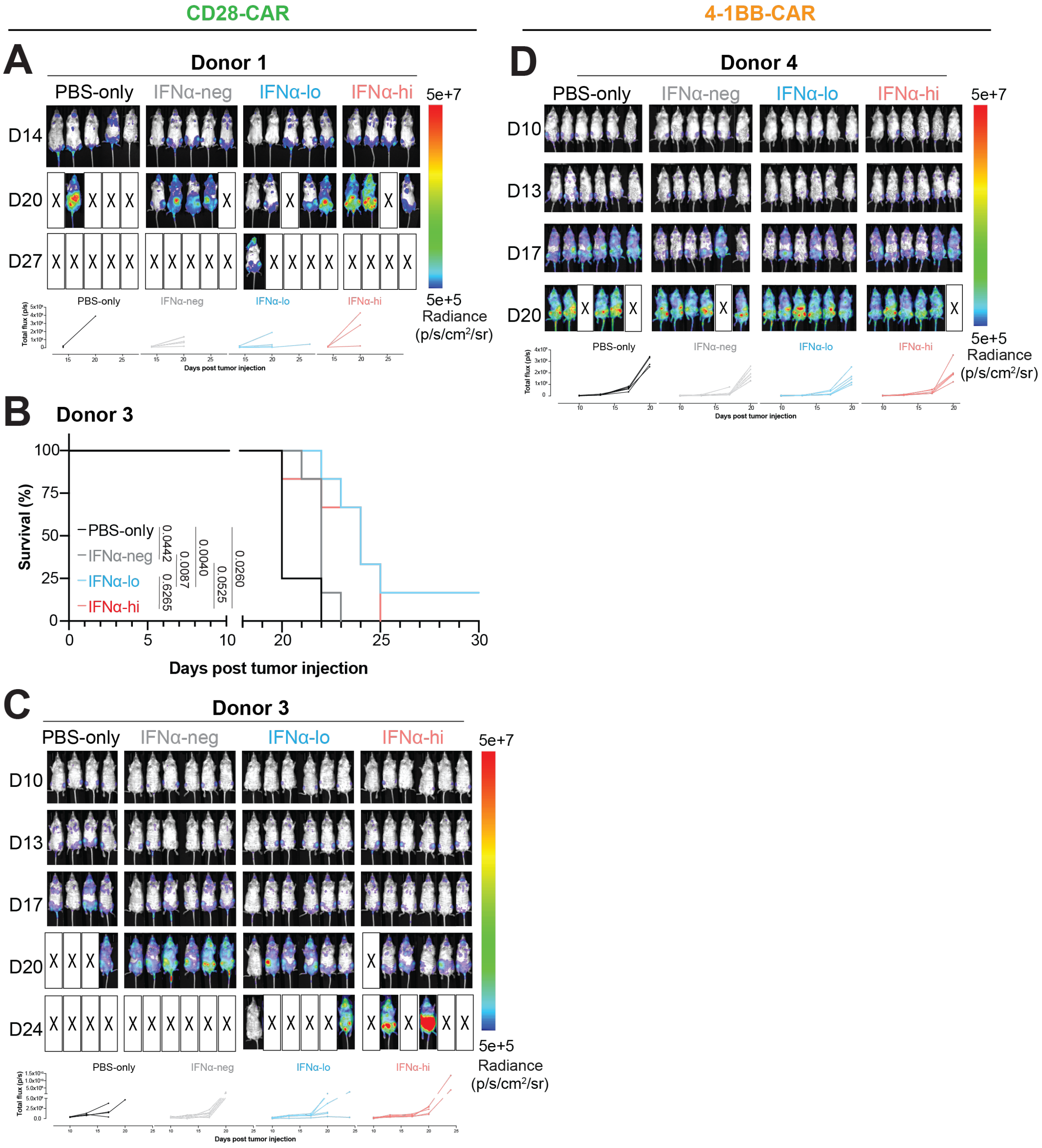
CD28-CAR
4-1BB-CAR
A
Donor 1
PBS-only
IFNα-neg
IFNα-lo
IFNα-hi
D14
D20
D27
5e+7
5e+5
Radiance
(p/s/cm²/sr)
Days post tumor injection
D
Donor 4
D10
D13
D17
D20
B
Donor 3
Survival (%)
PBS-only
IFNα-neg
IFNα-lo
IFNα-hi
0.0442
0.0087
0.0040
0.0260
0.6265
0.0525
Days post tumor injection
C
Donor 3
D24

**Figure S7. Survival of mice and tumor burden in OCI-Ly8 model following CAR T-cell engraftment.**

**A,** Representative bioluminescence imaging of tumor burden in mice treated with PBS or CD28-costimulated CAR T cells generated from donor 1 under the three IFNα culturing conditions (top), and corresponding quantification of bioluminescence signals (bottom, n=5,5,5,4). **B,** Kaplan-Meier analysis of survival of OCI-Ly8-bearing mice treated with PBS or $5\times10^5$ CD28-costimulated CAR T cells generated from donor 3 under the three IFNα culturing conditions (n=4,6,6,6). **C,** Serial bioluminescence imaging of tumor burden in the mice shown in (**B**, top), and corresponding quantification of bioluminescence signals (bottom). **D,** Serial bioluminescence imaging of tumor burden in mice treated with PBS or 4-1BB-costimulated CAR T cells generated from donor 4 under the three IFNα culturing conditions (top), and corresponding quantification of bioluminescence signals (bottom, n=6,6,6,6). Statistical analysis were performed by log-rank (Mantel-Cox) test.

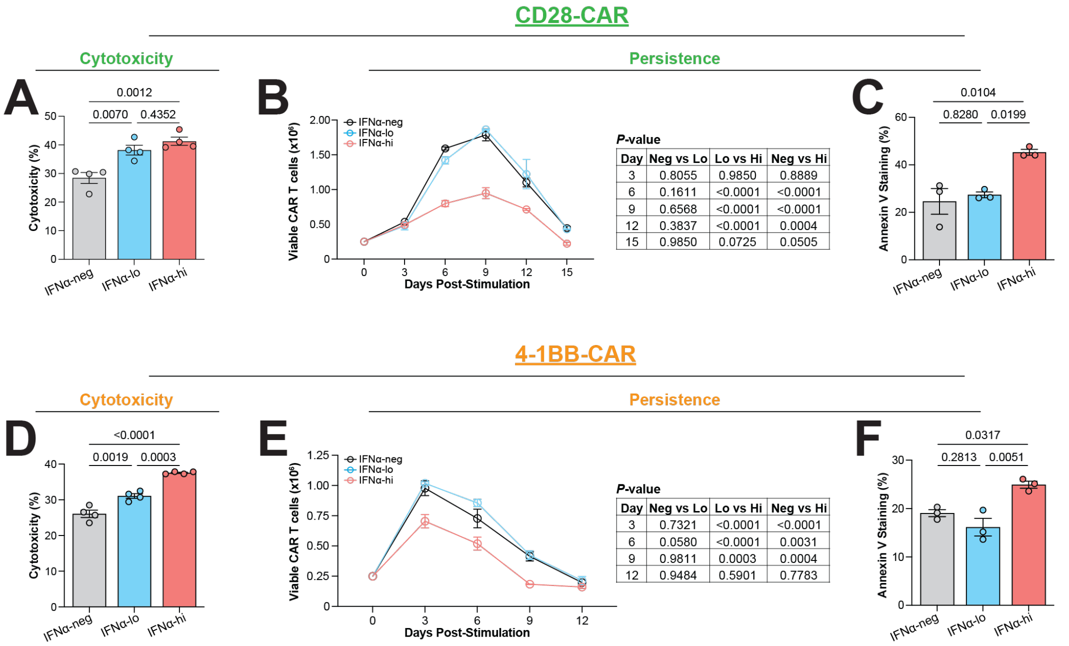


**Figure S8. CAR T-cell cytotoxicity on VAL cells and persistence after tumor stimulation.** **A** and **D,** Direct cytotoxicity of CD28-costimulated (**A**, n=3) and 4-1BB-costimulated (**D**, n=3) CAR T cells against firefly luciferase-expressing VAL target cells at E/T ratio of 1:1 via 6-hr bioluminescence assay. **B** and **E,** Line graphs depicting viable CD28-costimulated (**B**, n=3) and 4-1BB-costimulated (**E**, n=3) CAR T-cell cellularity in vitro following coculture with VAL at E/T ratio of 1:5. **C** and **F,** Apoptosis of CD28-costimulated (**C**, n=3) and 4-1BB-costimulated (**F**, n=3) CAR T cells from the three culturing conditions at the time points when cellularity declined in (**B**) and (**E**). Day 9 and day 6 after coculture, respectively. All data are shown as mean ± SEM of n experimental replicates from a representative donor. Results are representative of independent experiments using CAR T cells from 2 different donors. Statistical analysis were performed via one-way (**A**, **C**, **D**, and **F**) or two-way (**B** and **E**) ANOVA with Tukey's correction for multiple comparisons.

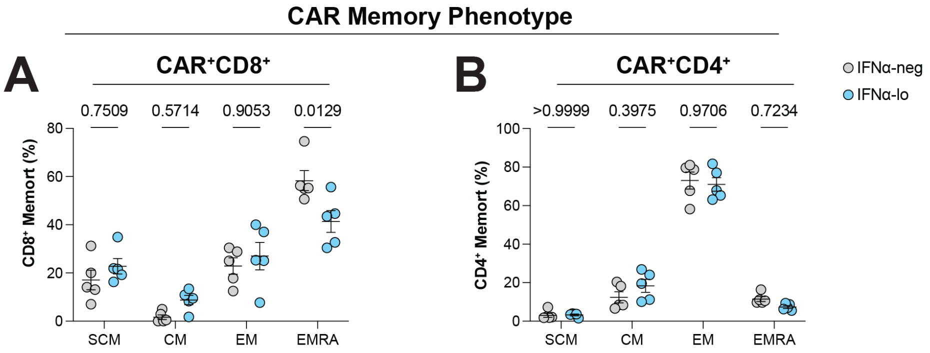


**Figure S9. Splenic CAR T-cell phenotype in the OCI-Ly8-based model for assessing durable CAR T-cell efficacy.**

**A** and **B,** Proportion of the SCM (CD45RA$^+$CD62L$^+$), CM (CD45RA$^-$CD62L$^+$), EM (CD45RA$^-$ CD62L$^-$), and TE (CD45RA$^+$ CD62L$^-$) memory phenotypes among CD8$^+$ and CD4$^+$ CAR T cells in spleen of mice shown in **Figure 6**. Statistical analysis were performed by two-way ANOVA with Tukey's correction for multiple comparisons.
